# An unconventional gatekeeper mutation sensitizes inositol hexakisphosphate kinases to an allosteric inhibitor

**DOI:** 10.1101/2023.04.26.538378

**Authors:** Tim Aguirre, Gillian L. Dornan, Sarah Hostachy, Martin Neuenschwander, Carola Seyffarth, Volker Haucke, Anja Schütz, Jens P. von Kries, Dorothea Fiedler

## Abstract

Inositol hexakisphosphate kinases (IP6Ks) are emerging as relevant pharmacological targets because a multitude of disease-related phenotypes has been associated with their function. While the development of potent IP6K inhibitors is gaining momentum, a pharmacological tool to distinguish the mammalian isozymes is still lacking. Here, we implemented an analog-sensitive approach for IP6Ks and performed a high-throughput screen to identify suitable lead compounds. The most promising hit, FMP-201300, exhibited high potency and selectivity towards the unique valine gatekeeper mutants of IP6K1 and IP6K2, compared to the respective wild-type kinases. Biochemical validation experiments revealed an allosteric mechanism of action that was corroborated by HDX-MS measurements. The latter analysis suggested that displacement of the *α*C helix, caused by the gatekeeper mutation, facilitates the binding of FMP-201300 to an allosteric pocket adjacent to the ATP binding site. FMP-201300 therefore serves as a valuable springboard for the further development of compounds that can selectively target the three mammalian IP6Ks; either as analog-sensitive kinase inhibitors or as an allosteric lead compound for the wild-type kinases.

## Introduction

Over the last decades, inositol pyrophosphates (PP-InsPs) have emerged as a group of ubiquitous small molecule messengers with pivotal functions in cellular homeostasis, metabolism and many disease-related signaling events.^[1]^ The densely charged molecules are interconverted in a fast metabolic cycle regulated by specific kinases and phosphatases (**Figure 1a**). Genetic knockouts of the three mammalian inositol hexakisphosphate kinase (IP6K) isozymes - enzymes involved in PP-InsP biosynthesis - have revealed their involvement in distinct biological processes, including insulin signaling^[2]^, apoptosis^[3]^, cancer metastasis^[4]^ and lifespan in mice.^[5]^ Despite catalyzing the same biochemical reaction, some IP6K functions were exclusively attributed to one isozyme. For example, knockdown of IP6K1, but not IP6K2, inhibited the exocytosis of insulin-containing granules in pancreatic beta cells.^[6]^ In another instance deletion of IP6K2, but not the other two IP6K isozymes, reduced cell death.^[7]^ Whether these observations are due to tissue-specific expression levels, localized cellular pools of PP-InsPs, or distinct modes of action by which the individual IP6Ks operate often remains elusive.

**Figure 1:**
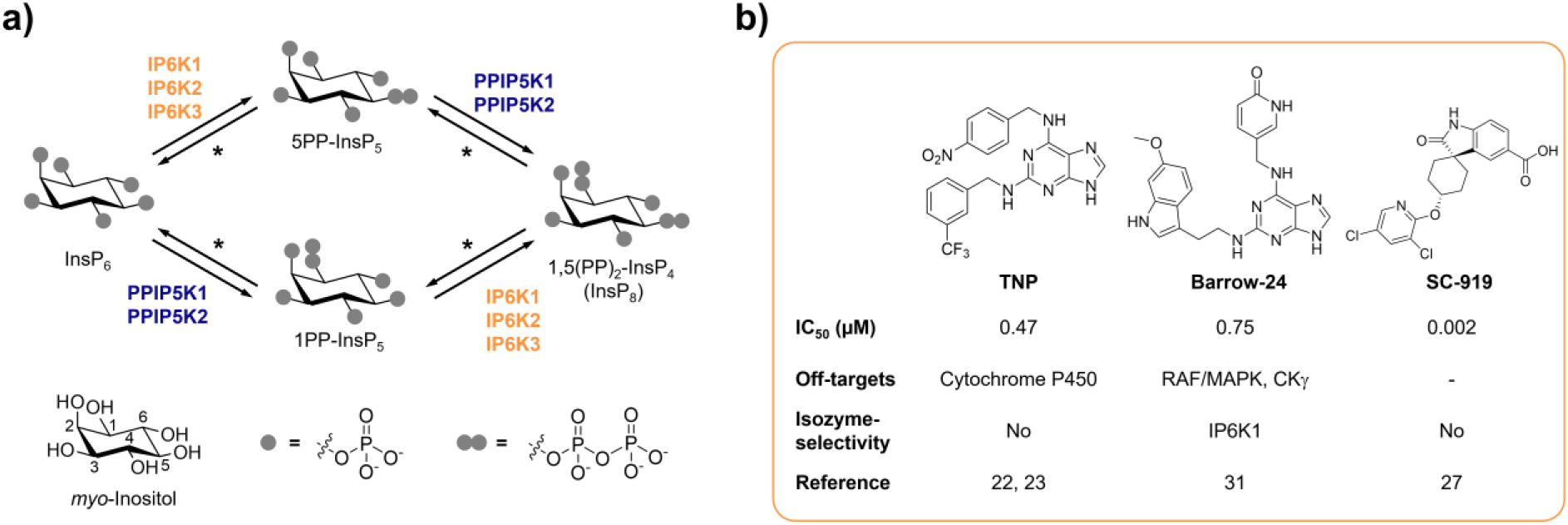
**a)** Simplified inositol pyrophosphate metabolism with focus on kinases. PPIP5K: Inositol hexakisphosphate and diphosphoinositol-pentakisphosphate kinase. *The PP-InsP dephosphorylation is catalyzed by nudix hydrolases DIPP1/2*α*/2*β*/3.^[18]^ **b)** Chemical structures and properties of selected IP6K inhibitors: widely used TNP, isozyme-selective Barrow-24, and most potent inhibitor SC-919.

Although genetic experiments perturbing IP6K function have contributed significantly to our understanding of PP-InsP signaling, these studies can be accompanied by compensatory effects and secondary genetic changes. The undesired transcriptional up- and downregulation of numerous genes has caused confounding results in the past.^[8]^ Furthermore, genetic knockout inevitably abolishes the non-catalytic functions of the kinase, which, under some circumstances, are essential for the regulation of specific cellular processes.^[3b, 9]^ The generation of single point kinase-dead mutants usually warrants correct protein expression and folding, but can be accompanied by notable transcriptional alterations.^[8]^ Lastly, a limitation of genetic approaches is the long timeframe, which can lead to genetic compensations^[10]^ and appears inherently incompatible with the rapid interconversion of PP-InsPs in cells, indicating fast signal transmission.^[11]^

A well-suited approach to address these challenges is the pharmacological inhibition of IP6K activity. While the benchmark inhibitor *N*2-(*m*-trifluorobenzyl)-*N*6-(*p*-nitrobenzyl)purine^[12]^ (TNP, **Figure 1b**) has lost appreciation due to the accumulation of unfavorable properties^[13]^, the development of novel pharmacological tools targeting IP6Ks has recently picked up speed.^[14]^ High-throughput screens and structure-guided design have yielded compounds with low nanomolar potencies, such as SC-919^[15]^ (**Figure 1b**) and others.^[16]^ However, with the exception of one isozyme-selective inhibitor targeting IP6K1^[17]^ (Barrow-24, **Figure 1b**), all other IP6K inhibitors lack specificity.

To achieve selective kinase inhibition, Shokat and co-workers developed the analog-sensitive approach, which enables the generation of mutually selective kinase-inhibitor pairs.^[19]^ A conserved medium-to-large sized gatekeeper residue is mutated to glycine or alanine, thereby creating an additional pocket in the ATP binding site that does not occur in any other wild-type (WT) kinase.^[19a, 20]^ While designed inhibitor analogs with a sterically demanding substituent fit into the enlarged ATP binding site, they clash with the active site of WT kinases.^[19a, 20-21]^ Implementation of this method for various kinase families of multiple organisms has provided remarkable insight into the function and regulation of phosphorylation-based signaling.^[22]^

While the analog-sensitive approach has been applied to over 80 protein kinases, the implementation for kinases phosphorylating small molecules, such as lipids or sugars, proved more challenging and has received scant attention to date. For example, mutation of the gatekeeper residue in *S. cerevisiae* VPS34, a PI3 kinase, yielded an active mutant allele, however, mutation of the mammalian PI3K p110*α* gatekeeper resulted in a drastic decline of kinase activity.^[23]^ A similar observation was made with PI3K-like kinase TOR2 from *S. cerevisiae*, which was sensitized to the mTOR inhibitor BEZ235 by mutating its gatekeeper residue to alanine. The mammalian kinase mTOR, on the other hand, was rendered catalytically inactive by the same mutation.^[24]^

Here, we implemented the analog-sensitive approach for inositol hexakisphosphate kinases to identify isozyme-selective inhibitors. While conventional alanine and glycine gatekeeper mutants rendered the kinase catalytically inactive, a leucine to valine mutant entirely retained its activity. A high-throughput screen with over 50.000 compounds yielded a potent and mutant-selective inhibitor of unknown biological activity. Validation and biochemical characterization revealed an allosteric mode of action, which was corroborated by HDX-MS measurements. This non-competitive mechanism of inhibition constitutes a promising new opportunity to selectively target the mammalian IP6K isozymes.

## Results

### Conventional gatekeeper mutations render IP6Ks catalytically inactive

The gatekeeper identification in inositol phosphate (InsP) kinases is facilitated by the structural similarity of their active site architecture to protein kinases.^[25]^ While the exact structures of mammalian IP6Ks have remained elusive to date, those of other members of the InsP kinase family, e.g. *Homo sapiens* IPMK and *Entamoeba histolytica* IP6KA, have successfully been deciphered.^[26]^ Owing to the decent sequence similarity within the active site, the gatekeeper residues of the human IP6K isozymes could be identified by simple sequence alignment with the abovementioned related InsP kinases and are Leu210, Leu206 and Leu201 for IP6K1, IP6K2 and IP6K3, respectively (**Figure 2a**).

**Figure 2:**
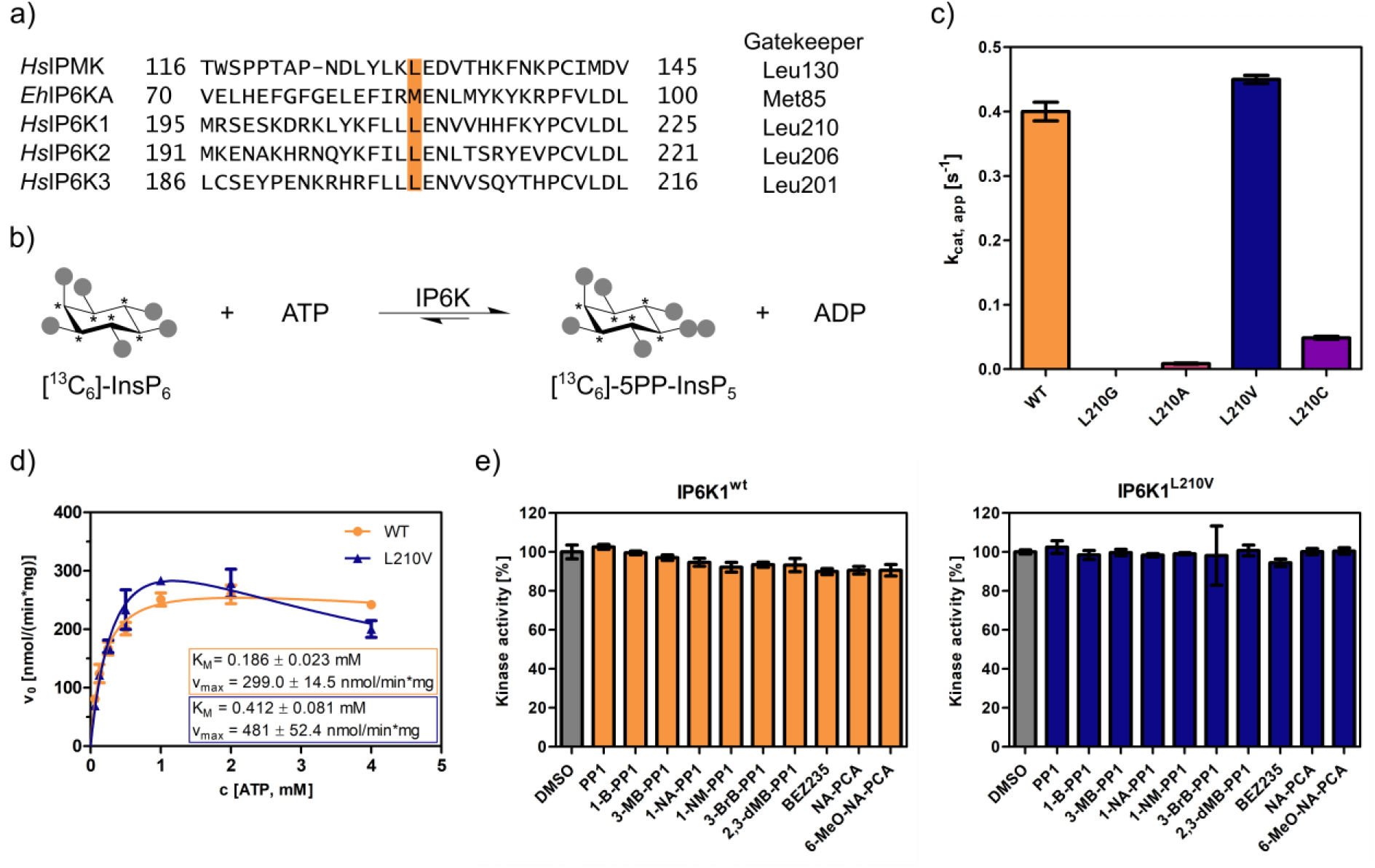
**a)** Sequence alignment of InsP kinases to identify the gatekeeper position of human IP6Ks. The gatekeeper residue is highlighted in orange. **b)** Kinase reaction of IP6Ks using uniformly ^13^C-labeled InsP_6_ as substrate. Asterisks indicate ^13^C-labeled positions. The conversion to 5PP-InsP_5_ was followed by an established spin-echo difference NMR method.^[27b, 29]^ **c)** Catalytic activity of IP6K gatekeeper mutants indicated by apparent turnover numbers k_cat, app_. **d)** Michaelis-Menten graphs for IP6K1^wt^ and IP6K1^L210V^. **e)** Screening of established analog-sensitive kinase inhibitors (**Figure S4**) at 10 μM concentration against IP6K1^wt^ and IP6K1^L210V^ using the NMR assay. All data points were measured in independent triplicates and error bars represent the standard deviation.

The gatekeeper mutants L210G and L210A of IP6K1 were generated by site-directed mutagenesis and the enzymes were recombinantly expressed, analogous to the WT kinase, as MBP-fusion proteins. To assess their catalytic activity, we used an NMR assay that harnesses fully ^13^C-labeled InsP6 as a substrate and enables direct product readout of 5PP-InsP5 (**Figure 2b and Figure S1**).^[27]^ Unfortunately, both the alanine and glycine mutations were not tolerated by IP6K1 and rendered the protein virtually inactive (**Figure 2c**). Likewise, IP6K2 suffered a drastic decline in substrate turnover upon gatekeeper mutation to glycine (**Figure S2**). While the cysteine mutant amenable for the electrophile-sensitive approach^[28]^ retained residual catalytic activity, established electrophile-sensitive kinase inhibitors did not inhibit the mutant kinase (**Figure S3**).

Since the size of the gatekeeper appeared to be relevant, the residue was subsequently mutated to valine; the last remaining hydrophobic amino acid larger than alanine but smaller than leucine. Although the change is comparably subtle, it could still have an impact on inhibitor selectivity. Unlike the previous mutations, the valine gatekeeper mutant completely retained its catalytic activity compared to the WT enzyme and was thus further investigated (**Figure 2c**).

### IP6K1^L210V^ is not susceptible to established analog-sensitive kinase inhibitors

To assess the impact of the valine gatekeeper mutation on the affinity of IP6K1 for ATP, Michaelis-Menten kinetics were measured for IP6K1^L210V^ and IP6K1^wt^ (**Figure 2d**). The KM value of IP6K1^wt^ was determined as 186 ± 23 μM and is in good accordance with the literature.^[27b, 30]^ The mutation of the gatekeeper residue did indeed impact the behavior of IP6K1 towards ATP, although the effects were relatively minor. We observed a two-fold reduction in ATP affinity (KM = 412 ± 81 μM) along with a 60% increase in vmax from 300 nmol*min^-1^*mg^-1^ to 480 nmol*min^-1^*mg^-1^, implicating a faster substrate conversion despite impaired ATP binding. The valine mutation also seems to exacerbate the substrate inhibition, which complicates the comparison of these steady-state parameters and suggests a change in protein dynamics upon gatekeeper mutation. Compared to many protein kinases, however, which often experienced a substantial reduction in ATP affinity upon gatekeeper mutation in the past^[31]^, the change of kinetic parameters between IP6K1^wt^ and IP6K1^L210V^ can be considered minor.

In a preliminary inhibitor screen, an array of established analog-sensitive kinase inhibitors was tested against both IP6K1^wt^ and IP6K1^L210V^. Among them were the most frequently used pyrazolopyrimidine (PP) analogs, BEZ235 and two 5-aminopyrazolo-4-carboxamide inhibitors (**Figure S4**). As expected, none of these compounds exhibited any inhibitory activity against IP6K1^wt^ at 10 μM concentration (**Figure 2e**). However, inhibition of IP6K1^L210V^ was not observed either, making these molecules unsuitable as inhibitors for analog-sensitive IP6Ks. Consequently, a novel inhibitor scaffold that allows for selective inhibition of analog-sensitive IP6Ks over their WT counterparts needed to be identified.

### Luminescence-based assay of reverse kinase reaction enables pilot screen

Seeking to identify an inhibitor selective for the gatekeeper mutant, IP6K1^wt^ and IP6K1^L210V^ were submitted to a high-throughput screen encompassing 54.912 small molecule inhibitors and inhibitor-like structures.^[32]^ Although the NMR assay described above provides direct product detection and the ability to distinguish the PP-InsP isomers^[27b]^, it is inherently slow and thus not compatible with a high-throughput screen. The use of luminescence-based readouts of ATP consumption to measure IP6K activity is error-prone due to the kinase’s unusually low affinity for ATP, the sensitivity to the ATP/ADP ratio, and its inherent ATPase activity.^[30b, 33]^ Nevertheless, we were able to take advantage of these properties by examining the reverse reaction of IP6K1 with the Kinase-Glo^®^ assay, where 5PP-InsP5 and ADP are converted to InsP6 and ATP (**Figure 3a**). Hence, physiologically relevant nucleotide concentrations could be used without causing ambiguity, since the amount of generated ATP was directly dependent on the substrate concentration. Similar KM values for ATP and ADP should allow for comparability of inhibitor potencies,^[33]^ and a gain-of-signal detection (rather than loss-of-signal) resulted in a much better signal-to-noise contrast in our setting.

**Figure 3:**
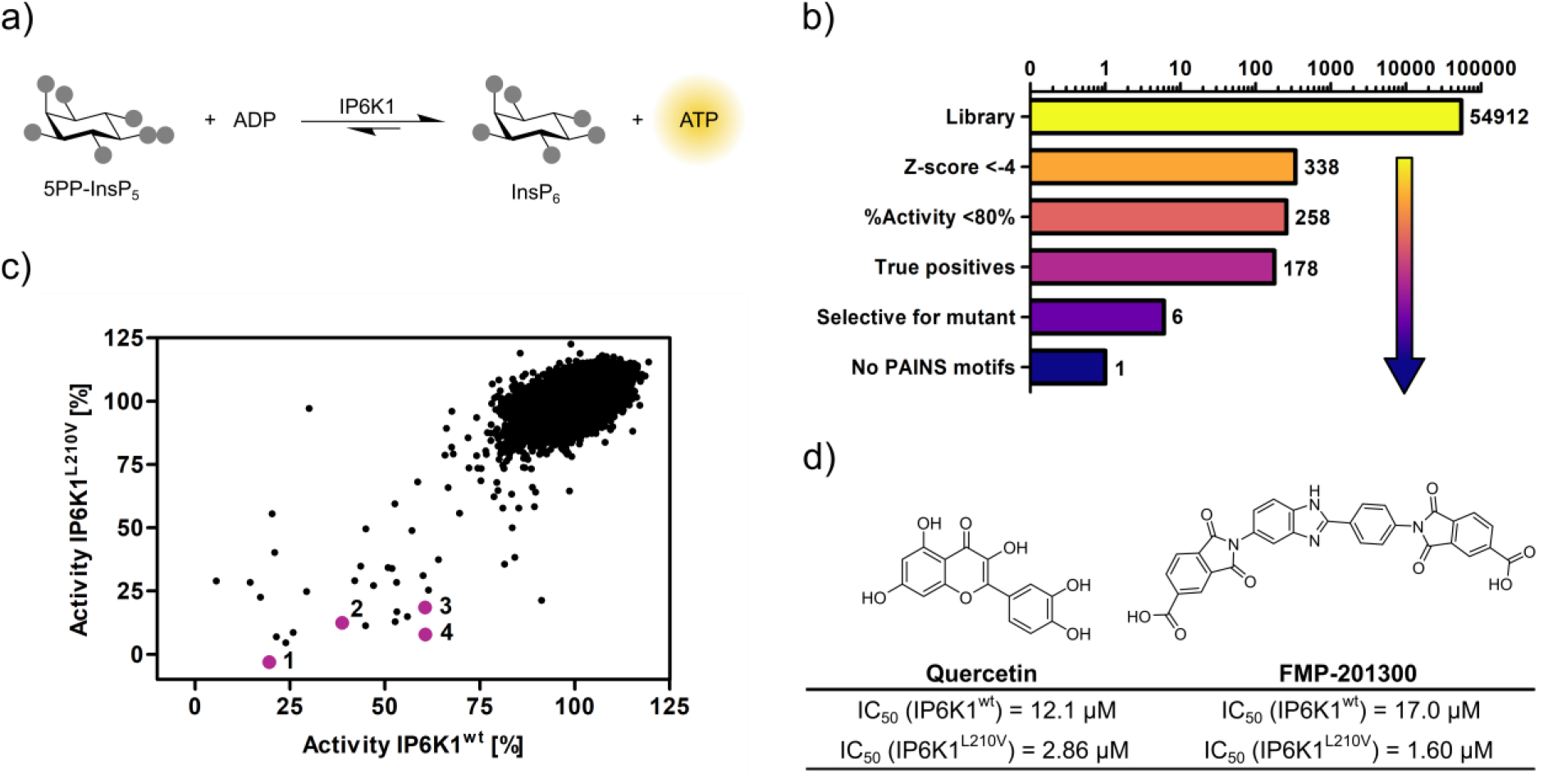
**a)** ATP synthase reaction used for the high-throughput screen. The generation of ATP was monitored by the luminescence-based Kinase-Glo^®^ assay. **b)** Reduction of putative hits by application of specific thresholds and manual selection. **c)** Scatter plot comparing the potency of compounds measured in the primary screen against IP6K1^wt^ and IP6K1^L210V^. Hits highlighted in magenta represent known IP6K inhibitors, or promising hits. 1: 6-hydroxy-dl-DOPA, 2: FMP-201300, 3: myricetin, 4: quercetin. **d)** Examples of true positive hits that display selectivity for IP6K1^L210V^, including known IP6K inhibitor quercetin and newly discovered FMP-201300.

After optimizing reaction parameters to ensure a successful and robust high-throughput screen (**Figure S5a**,**b**), an IC50 curve of the known IP6K inhibitor myricetin was recorded to determine the accuracy of the reverse reaction (**Figure S5d**).^[30a, 34]^ The measured value of 3.1 μM is in the same range as the published value for the forward reaction^[30a]^, thereby strengthening the initial rationale. To further confirm the assay viability, a Z’-plate was measured employing positive and negative controls only (**Figure S6a**). An excellent Z’-factor^[35]^ of 0.81 was achieved indicating high assay quality and feasibility. Before screening the entire library, a pilot screen encompassing 1670 FDA-approved drugs and 1280 pharmacologically active compounds was conducted. It confirmed the suitability of the final screening setup by yielding known IP6K inhibitors myricetin and 6-hydroxy-DL-DOPA^[30a, 34]^ and displaying good hit reproducibility as depicted in the Bland-Altmann plot (**Figure S6b**).^[36]^

### High-throughput screening identifies a mutant-selective inhibitor

The successful pilot screen was followed by screening 54.912 compounds comprising diverse chemical motifs^[32]^ and natural products against IP6K1^L210V^ and IP6K1^wt^. Putative hits for the gatekeeper mutant were subsequently identified by excluding compounds with a z-score higher than –4 and a residual activity greater than 80% (**Figure 3b**). The remaining 258 compounds (0.5% hit rate) were retested in a dose-dependent reconfirmation and counter screen (see SI for details) to eliminate random false positives (compounds that do not confirm activity) and chemical false positives (compounds directly interfering with the assay readout). Additionally, information was added on compounds identified as frequent hitters in our in-house screening data. This analysis uncovered 80 molecules as false positives, while 178 inhibitors were consequently classified true hits. However, only a handful appeared to be selective for IP6K1^L210V^ as evident from the scatter plot comparing the residual activities of both kinases (**Figure 3c**). Curiously, the majority of mutant-selective inhibitors were dietary flavonoids such as quercetin and fisetin (**Table S1**).^[34]^

Since flavonoids are promiscuous ATP-competitive inhibitors with various off-target effects^[37]^, and the compounds had been flagged as pan-assay interference compounds (PAINS) based on structural filters^[38]^, they were deemed unsuitable as selective IP6K inhibitors and thus not investigated further. The only non-polyphenolic molecule, FMP-201300, displayed good selectivity for the gatekeeper mutant with IC50 values of 17 μM and 1.6 μM against IP6K1^wt^ and IP6K1^L210V^, respectively (**Figure 3d**). Importantly, the almost symmetrical inhibitor has no known inhibitory activities against other biological targets and does not display any recognized PAINS motifs.^[38]^ Furthermore, FMP-201300 was neither redox-active, nor cytotoxic, against HEK293 or HepG2 cell lines at a concentration of 10 μM (**Tables S3 and S4**).

### Potency of FMP-201300 is dictated by carboxylic acid functional groups

Next, FMP-201300 was validated with the NMR assay using isotopically labeled [^13^C6]-InsP6 as a substrate, and 1 mM ATP, as well as a creatine kinase/phosphocreatine-based ATP-regenerating system. The assay confirmed FMP-201300 as a potent and mutant-selective inhibitor with IC50 values of 114 nM and 810 nM for IP6K1^L210V^ and IP6K1^wt^, respectively (**Figure 4a**). To evaluate whether the inhibitory behavior of FMP-201300 was transferable to the gatekeeper mutant of another IP6K isozyme, dose response curves against IP6K2^wt^ and IP6K2^L206V^ were recorded. FMP-201300 was 11 time more potent against IP6K2^L206V^ compared to IP6K2^wt^ with IC50 values of 190 nM and 2.14 μM, respectively (**Figure 4b**). Due to difficulties in obtaining recombinant catalytically active IP6K3^wt^, this isozyme could not be investigated. However, the evolutionarily related ortholog IP6KA from *Entamoeba histolytica* (*Eh*) was sensitized to inhibition by FMP-201300 when its methionine gatekeeper was mutated to valine (**Figure S7a**). The approach therefore appears transferable not only between the mammalian isozymes, but also to other IP6K orthologs.

**Figure 4:**
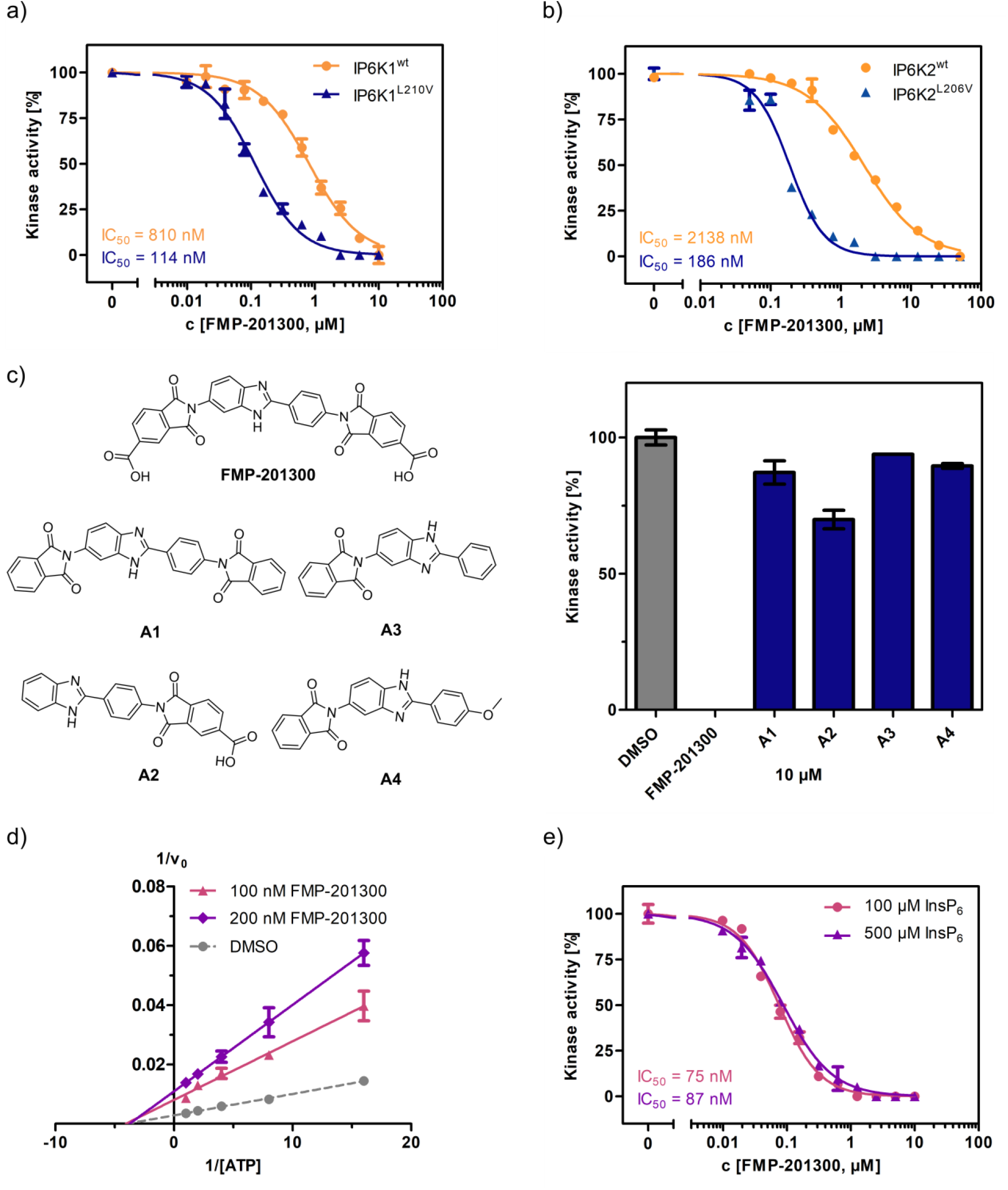
**a)** IC_50_ curves of FMP-201300 against IP6K1^wt^ and IP6K1^L210V^. **b)** IC_50_ curves of FMP-201300 against IP6K2^wt^ and IP6K2^L210V^. 100% activity corresponds to the DMSO control and 0% indicates no substrate conversion. **c)** Inhibitory activity of FMP-201300 analogs against IP6K1^L210V^ at 10 μM concentration. **d)** Lineweaver-Burk plot of FMP-201300 against IP6K1^L210V^ at two different inhibitor concentrations. The plotted lines for the DMSO control and two different inhibitor concentrations intersect almost precisely on the x-axis, indicating no change in K_M_ value and a decrease in v_max_ upon inhibition **e)** IC_50_ curves of FMP-201300 against IP6K1^L210V^ at two different InsP_6_ concentrations. All data points were measured in independent triplicates and error bars represent the standard deviation.

Given these promising initial results, we wanted to understand which elements of the peculiar structure of FMP-201300 are potentially involved in steering inhibitory activity towards the gatekeeper mutant. Therefore, the MolPort chemical library was screened for compounds containing a structure similar to FMP-201300 or core substructures such as phthalimides or benzimidazoles. Four analogs (**A1**-**A4**) were subsequently tested against IP6K1^L210V^ at a single concentration of 10 μM (**Figure 4c**). Interestingly, none of the analogs inhibited the kinase by more than 30%. Although **A1** was merely lacking the carboxylic acid groups, it was completely inactive. The only analog still possessing a carboxylic acid moiety, **A2**, inhibited IP6K1^L210V^, albeit with severely reduced potency. This result is consistent with the observation made with IP6K inhibitors SC-919^[15b]^ and LI-2172^[16b]^, which share the benzoic acid substructure with FMP-201300. Hence, the carboxylic acid appears to be indispensable for inhibitory activity and is likely engaged in pivotal interactions at the inhibitor binding site.

### Kinetic characterization reveals an allosteric mode of inhibition

Before undertaking further analyses to elucidate the structural determinants of mutant selectivity, we wanted to confirm the ATP-competitive inhibition mechanism of FMP-201300. Therefore, Michaelis-Menten measurements in the presence of inhibitor were performed and plotted reciprocally to obtain Lineweaver-Burk graphs.^[39]^ Surprisingly, these measurements revealed a clear allosteric mechanism for FMP-201300 (**Figure 4d**). To dispel any doubts about the accuracy of the Lineweaver-Burk plots, the assay was performed on the established IP6K inhibitor TNP. In agreement with the literature, the results indicated an unambiguous competitive mechanism of action for TNP (**Figure S8a**). In addition, TNP exhibited no significant difference in potency against IP6K1^wt^ and IP6K1^L210V^, again pointing towards a different inhibitory mechanism of FMP-201300 (**Figure S8b**).

These results suggest that the leucine to valine gatekeeper mutation – despite being buried deep in the ATP-binding pocket – must have caused a conformational change within the kinase structure that facilitates binding of FMP-201300. The fact that the inhibitor exhibits the same behavior against IP6K2 and *Eh*IP6KA diminishes the likelihood of a distant allosteric site, since IP6K1 and IP6K2 only share 47% sequence identity, most of which lies within the catalytic region. Since a structural alteration at a vastly remote site of IP6K1, caused by the subtle amino acid change, appears unlikely, we hypothesize that the compound binds adjacent to the ATP binding site.

Although FMP-201300 does not resemble the densely charged InsP6 molecule, the two carboxylic acids could potentially contribute to a substrate-competitive mechanism. Due to the nanomolar KM of IP6K1 for InsP6[33] and the inherently low sensitivity of the NMR assay, Michaelis-Menten measurements were not feasible. The Cheng-Prusoff equation states that for [S] >> KM, the IC50 values correlate linearly with substrate concentration for competitive inhibitors.^[40]^ However, the difference in IC50 values measured at 100 μM and 500 μM InsP6 concentration was negligible, excluding substrate inhibition as a potential mechanism (**Figure 4e**).

### HDX-MS corroborates allosteric mechanism

For a more detailed picture of the specific binding modalities of FMP-201300 and the factors promoting selectivity for the gatekeeper mutant, co-crystal structures of inhibitor bound to the kinase would undoubtedly be beneficial. However, to date the mammalian IP6K isozymes have been recalcitrant in crystallization efforts. This limitation may, in part, be circumvented by focusing on *Eh*IP6KA, which has been crystallized and structurally analyzed.^[26b]^ An alignment of the AlphaFold structure model of IP6K1 and the crystal structure of *Eh*IP6KA suggests significant structural conservation, especially within the active site (**Figure S7c**). The fact that FMP-201300 recapitulates its gatekeeper mutant-selectivity and allosteric binding mode against *Eh*IP6KA^M85V^ (**Figure S7a**,**b**) prompted us to attempt co-crystallization of this related kinase and the inhibitor. While the valine gatekeeper mutant of *Eh*IP6KA was readily crystallized using the published conditions for the WT counterpart^[26b]^, soaking or co-crystallization with FMP-201300 remained unsuccessful. Remarkably, the structures of the WT and mutant *Eh*IP6KA are virtually identical except for the additional space generated by the reduced gatekeeper size (**Figure S7d**). This result illustrates a potential limitation of crystallography, as it can only provide a snapshot of a momentary protein conformation and may not be suited to unveil subtle dynamic changes that occur upon inhibitor binding.

We therefore utilized hydrogen deuterium exchange mass spectrometry (HDX-MS) to elucidate which parts of the protein participate in inhibitor binding. Initially, we compared the deuterium exchange rates between the IP6K1^L210V^ apo and IP6K1^L210V^ bound to FMP-201300. Major decreases in exchange rates were observed in proximity to the ATP binding site, whereas remote sites were mostly unaltered (**Figure 5A and Figure S9a**). Amino acid residues 40-45, 67-79 and 205-212, corresponding to *β* strands 1-3, and residues 53-58 within the *α*C helix of IP6K1, exhibited large decreases in deuterium exchange rates. Importantly, this includes the gatekeeper residue Leu210 and parts of the hinge region (Glu211-Asn212-Val213), corroborating the hypothesis of ATP-adjacent binding. In agreement with previous kinetic measurements that suggested a non-competitive mechanism, changes in ATP binding regions were less pronounced with no differences that met the criteria for significance for the ATP ribose-binding residue Asp224 (**Figure S9**).

**Figure 5:**
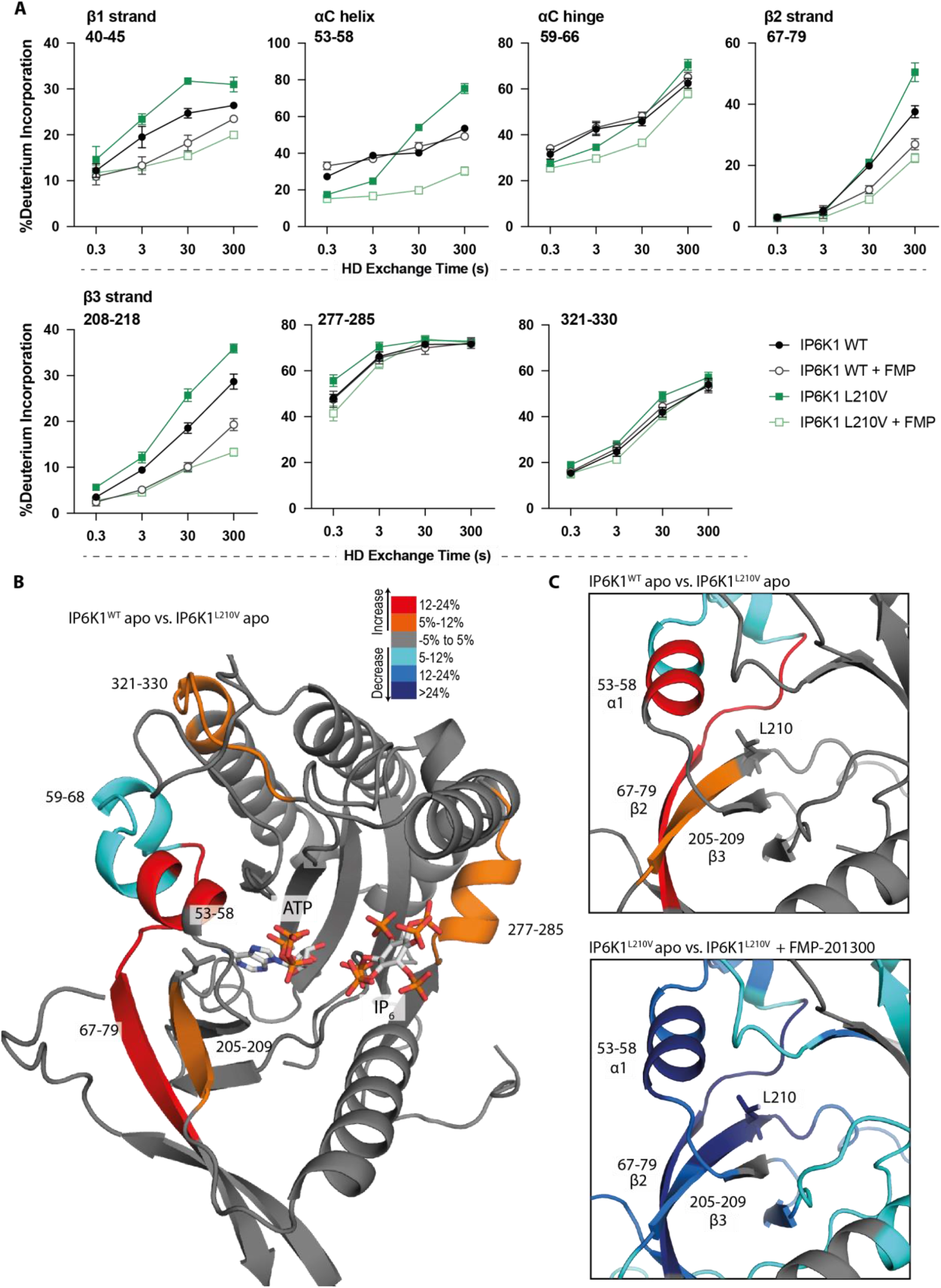
HDX-MS reveals increased flexibility in the *β*2-*β*3 strands and *α*C helix induced by the IP6K1^L210V^ mutation that sensitize the enzyme to FMP-201300. **A:** HDX differences in IP6K1. Time course of deuterium incorporation for a selection of peptides. Raw data can be found in the supplemental file. **B:** Overall HDX-MS changes in deuterium incorporation induced by the IP6K1^L210V^ mutation. Differences in deuterium exchange rates mapped on a model of IP6K1^wt^ (Alpha-Fold structure prediction Q92551 with ATP and InsP_6_ from *Eh*IP6KA docked in the active site (PDB: 4O4F)). Peptides that showed significant differences in HDX and met the cut-offs were included (>6% deuterium incorporation and 0.5 Da with an unpaired student t-test of p<0.05). Strongly disordered regions and regions of low per-residue confidence scores (pLDDT) were omitted for clarity. **C**: A magnified view of gatekeeper region included the *β*2-*β*3 strands and *α*C helix. The same regions that show increased flexibility in the IP6K1^L210V^ mutant (top panel) also undergo large decreases in exchange upon binding to FMP-201300 (bottom panel). This increased flexibility likely allows accommodation of the inhibitor and ATP.

Decreases in HD exchange could be indicative of direct interactions of the inhibitor with IP6K1, which would support an allosteric mode of action. Specifically, FMP-201300 might bind in a cleft next to the ATP site, which stretches to *β*1-*β*3 strands and *α*C helix, while engaging with the gatekeeper residue. Similar binding modes have been observed for other allosteric regulators, including EGFR and MEK inhibitors.^[41]^ Interestingly, crucial structural features of IP6K1 such as the IP helices and the PDKG catalytic motif did not exhibit differences in deuterium exchange rates, indicating no direct binding interaction or induced structural changes at these sites (**Figure S9**). Conversely, the conserved IDF tripeptide (Ile398-Asp399-Phe400) was moderately affected by binding of FMP-201300, potentially influencing ATP binding as Asp399 coordinates two Mg^2+^ ions and makes polar contacts with the nucleotide phosphates. Furthermore, a unique 310A helix comprising residues 278-284 showed a medium difference in deuterium exchange rates. This helix is located opposite the IP helices in the substrate-binding pocket and makes pivotal interactions with InsP6 (**Figure S9**). Since previous biochemical experiments excluded substrate competition of FMP-201300, this region likely undergoes dynamic structural changes rather than being involved in direct binding. Nevertheless, these structural changes could conceivably affect the kinetic properties of IP6K1, as residues Lys278 and Arg282 are involved in the positioning of InsP6. Hence, a change in their orientation could impact InsP6 binding and thereby kinase catalytic activity.

To understand the impact of the gatekeeper mutation on protein conformation, we next investigated the differences in deuterium exchange rates between the IP6K1^wt^ and IP6K1^L210V^ apo forms. The leucine to valine mutation led to increases in HD exchange in the *β*2-*β*3 strands and *α*C helix, indicating an overall increased flexibility in this region (**Figure 5**). A minor decrease in exchange was also identified at a flexible region immediately adjacent to the *α*C helix (residues 59-68) that flanks the active site and is a dynamic modulator of kinase activity.^[42]^ Despite the minor differences between the structures of the gatekeeper residues, the leucine appears to be important for the stability of these features, although it is difficult to model without a high-resolution structure of the human enzyme. The associated structural rearrangements induced by the gatekeeper mutation could be responsible for the mutant-selectivity of FMP-201300, since targeting the displacement of *α*C helices has been a viable approach for the development of several allosteric inhibitors.^[43]^ For instance, the T790M resistance gatekeeper mutation in epidermal growth factor receptor (EGFR) sensitized the tyrosine kinase to allosteric inhibitors by an outward displacement of the *α*C helix in the inactive conformation.^[41a]^ Small increases in deuterium uptake also occurred in a flexible loop adjacent to the *α*C helix and a distant helix that could be important for substrate binding (residues 321-330 and 277-285), indicating further increased flexibility of the secondary structure in the mutant enzyme that allows for the accommodation of FMP-201300, ATP, and substrate (**Figure 5**).

Finally, we wondered whether FMP-201300 would impose similar changes in deuterium uptake on IP6K1^wt^. We therefore compared the apo kinase to the inhibitor-bound form. With the exception of the *β*2 and *β*3 strands and parts of the hinge region that displayed a slight decrease in HDX, no significant changes in deuterium incorporation were observed (**Figure S9a**). These data underline the selectivity and preferred binding of FMP-201300 to IP6K1^L210V^. In conclusion, the L210V gatekeeper mutation of IP6Ks putatively causes a displacement of the *α*C helix sufficient to expand a pocket adjacent to the ATP binding site, which sensitizes the kinase to an allosteric inhibitor. Alternatively, the mutation induces a certain degree of flexibility that facilitates the induced-fit binding and *α*C helix displacement by FMP-201300.

## Discussion

We have investigated the suitability of an analog-sensitive approach for the selective inhibition of the mammalian IP6K isozymes. While the conventional glycine and alanine gatekeeper mutants suffered a drastic decline in catalytic efficiency, a subtle leucine to valine mutation did not perturb the kinase activity. After established analog-sensitive kinase inhibitors failed to target IP6K1^L210V^, a high-throughput screen uncovered FMP-201300 as a potent and mutant-selective inhibitor. The compound had not been assigned any function in previous screens and displayed no toxicity in cell culture. A preliminary structure-activity analysis indicated the indispensability of the carboxylic acid as a structural motif required for inhibitory potency. This phenomenon had been observed in previous studies investigating IP6K inhibitors, where the removal of a carboxylate moiety from the core structure resulted in a drastic decrease in potency of up to 2600-fold.^[16]^ While this functional group appears essential for inhibitory activity, it can pose several drawbacks such as metabolic instability, toxicity, and poor passive diffusion across biological membranes. To assess cell permeability and cellular activity in a preliminary experiment, we measured the reduction of ribosomal RNA synthesis upon treatment with FMP-201300 (**Figure S10**). The inhibitor, alongside two established IP6K inhibitors, was able to recapitulate this recently reported phenotype^[44]^, indicating sufficient crossing of the cell membrane despite its two carboxylic acids. In fact, other carboxylate-containing IP6K inhibitors had proven efficacious *in cellula* and *in vivo* before.^[15a, 16a]^ Regardless, a wide range of bioisosteres, including hydroxamic acids, sulfonamides, tetrazoles, and others^[45]^, would be available to circumvent potential issues arising with FMP-201300.

Unexpectedly, Lineweaver-Burk measurements suggested an allosteric mechanism of FMP-201300, which appeared counterintuitive with regards to the analog-sensitive approach. After an InsP6-competitive mode of action had been ruled out, attempts to gain structural insights by crystallographic analysis of the ortholog *Eh*IP6KA were undertaken. Although the valine gatekeeper mutant of *Eh*IP6KA was successfully crystallized, soaking or co-crystallization of FMP-201300 could not be achieved. Lastly, HDX-MS measurements were performed to obtain information on the structural determinants of mutant-selectivity. This analysis revealed distinct interaction profiles of FMP-201300 with the WT and mutant kinase and suggested an *α*C helix displacement to be responsible for sensitizing IP6K1 to allosteric inhibition.

The reduction of catalytic activity upon introduction of a space-creating gatekeeper mutation is by no means an issue exclusively occurring in IP6Ks. In fact, a loss in activity was observed for roughly 30% of all examined protein kinases^[46]^ and has proven particularly detrimental for small molecule kinases, such as the phosphoinositide lipid kinases.^[23-24]^ To address this challenge, Shokat and co-workers identified second-site suppressor mutations that rescue the catalytic activity of some kinases sensitive to the reduction in gatekeeper size.^[46]^ Since IP6Ks possess a protein kinase fold^[25]^, they might be amenable to second-site suppressor mutations as well.

As an alternative, we implemented a valine gatekeeper mutation that was well tolerated by IP6Ks. This unconventional approach could, in fact, constitute a viable alternative for kinases that are incompatible with the classical gatekeeper mutations. While the substitution of leucine with valine was comparatively subtle, the majority of kinases in the human kinome carry medium- to large-sized gatekeeper residues, such as methionine, tyrosine, or phenylalanine.^[47]^

Although the analog-sensitive method is a very promising tool, the prospect of having FMP-201300, or analogs thereof, as well-defined allosteric inhibitors is also very useful. The greater heterogeneity of residues and conformations in allosteric pockets^[48]^, even in isozymes of the same kinase, could pave the way for isozyme-selective IP6K inhibitors that do not require chemical genetic engineering. However, the kinetic measurements and HDX-MS analysis are insufficient to confirm the exact binding position with certainty, necessitating further biophysical measurements such as limited proteolysis MS^[49]^, cryo-EM^[50]^, or NMR using ^15^N-labeled protein.

Binding of FMP-201300 to a hydrophobic pocket adjacent to the ATP site, which is generated by *α*C helix displacement, seems plausible and is an established mode of action for allosteric kinase inhibitors.^[43]^ It remains unclear if the gatekeeper mutation caused this displacement or merely increased the conformational flexibility of the *α*C helix to facilitate the induced fit of FMP-201300. The first scenario appears more likely, as a similar phenomenon was observed for an allosteric EGFR inhibitor that was 1000-fold more potent against the acquired T790M gatekeeper resistance mutant, compared to the WT kinase.^[41a, 51]^ The EGFR inhibitor also exhibited direct binding interaction with the mutated gatekeeper residue, similar to our observations with FMP-201300. In general, these allosteric inhibitors preferably target an inactive conformation of protein kinases, which often confers selectivity owing to their greater structural heterogeneity compared to the corresponding active conformations.^[52]^ It could therefore be worthwhile to investigate and allosterically target the inactive conformation of IP6Ks.

The biochemical characterization of FMP-201300 has taken a surprising yet fascinating turn that necessitates follow-up investigations. It needs to be assessed whether the structural change putatively caused by the gatekeeper mutations has an impact on known protein-protein interactions and scaffolding functions of IP6Ks. Furthermore, it will be interesting to see if this allosteric pocket can be efficiently targeted without prior mutation of the gatekeeper. Lastly, a suitable cellular phenotype to study the inhibitor’s efficacy and specificity in a biological setting would be highly desirable. As of now, FMP-201300 constitutes a valuable addition to the ever-growing group of IP6K inhibitors and serves as an intriguing starting point for further characterization and improvement. It could provide a springboard for the heavily sought-after isozyme-selective inhibition of mammalian IP6Ks, either as analog-sensitive kinase inhibitor or allosteric lead compound for the development of inhibitors against the wild-type kinases.

## Methods

### Ligation-independent cloning and PCR mutagenesis

The mammalian IP6K ORFs were subcloned from a pTrc-His vector into a pET-His6-MBP-N10-TEV vector by ligation-independent cloning using the primers listed in **Table S5**. Single point mutations were subsequently installed by site-directed mutagenesis PCR using a standard protocol and the primers listed in **Table S6**. Detailed procedures can be found in the *Supporting Information*.

### Expression and purification of MBP-IP6K constructs

MBP-IP6K constructs were transformed into *E. coli arctic express* cells and expressed at 13°C overnight. Purification was performed using immobilized metal ion affinity chromatography, anion exchange chromatography and size-exclusion chromatography in succession. Detailed procedures can be found in the *Supporting Information*.

### Expression and purification of *Eh*IP6KA constructs

*Eh*IP6KA was expressed like described before^[27b]^ with slight alterations. In brief, *Eh*IP6KA^wt^ and gatekeeper mutant constructs were expressed in *E. coli arctic express* at 13°C overnight. The cells were lysed with a microfluidizer™ LM10 at 15000 psi and the debris removed by centrifugation. Purification was performed using immobilized metal ion affinity chromatography with imidazole elution. Detailed procedures can be found in the *Supporting Information*.

### NMR activity assays and IC50 determination

NMR assay for IP6K activity was performed similarly like described before^[27b]^ with slight alterations. In brief, reactions were run in 150 μL total volume containing 20 mM buffer, 50 mM NaCl, 6 mM MgCl2, 1 mM ATP, 0.2 mg/mL BSA, 1 mM DTT, 5 mM creatine phosphate and 1 U/mL creatine kinase (ATP-regenerating system), 100 μM InsP6 and 50 nM kinase in D2O if not otherwise stated. If applicable, inhibitor was added as a DMSO-*d6* stock in a two-fold dilution series at a final DMSO concentration of 1%. The reaction was run at 37°C and quenched with 400 μL quenching solution (20 mM EDTA pH* 6.0, 68.75 mM NaCl, in D2O) before being analyzed *via* NMR. Detailed procedures can be found in the *Supporting Information*.

### Kinase-Glo^®^ assay optimization and high-throughput screen

The reactions were carried out in 384-well plates. 5 μL 4× kinase buffer (buffer only for no kinase controls), 5 μL 4× DMSO or inhibitor and 5 μL 4× ADP solution were subsequently added to all wells. After 10 minutes equilibration, 5 μL 4× 5PP-InsP5 were added to start the reaction (milli-Q water for negative controls). After three hours at room temperature, 20 μL of *Promega* Kinase-Glo Plus^®^ reagent were added and the luminescence read out after 10 minutes of equilibration. For the high-throughput screen, ADP was added as 2× stock solution (10 μL) and compounds as 1 mM DMSO stock (0.2 μL). Detailed procedures can be found in the *Supporting Information*.

### Michaelis-Menten and Lineweaver-Burk kinetics

Kinetic assays were performed in D2O in a total volume of 500 μL containing the same buffer components for IP6K and *Eh*IP6KA constructs like stated for the NMR activity assays above. However, only 5 mM MgCl2 were added as a constant amount while ATP was added as ATP*Mg solution prepared in two-fold dilution starting from 50 mM. ATP*Mg was added to final ATP concentrations ranging from 4 mM to 62.6 μM. If applicable, inhibitor was added as a DMSO-*d6* stock at a final DMSO concentration of 1%. The reaction was run at 37°C and quenched with 38 μL 700 mM EDTA pH* 8.0 in D2O before being analyzed *via* NMR. Detailed procedures can be found in the *Supporting Information*.

### Crystallization of *Eh*IP6KA^M85V^ (27-270)

The construct for crystallization was expressed similarly to the construct for biochemical characterization with slight alterations. In brief, the kinase was expressed in *E. coli arctic express* from pDest-566-MBP-*Eh*IP6KA^M85V^ (27-270) at 13°C overnight. The protein was purified by immobilized metal ion affinity chromatography and the MBP-tag was subsequently removed by treating with TEV protease overnight.

For crystallization, the purified *Eh*IP6KA^M85V^ variant protein was complexed with 10 mM MgCl2 and 10 mM ATP, and crystallized using the sitting-drop vapor-diffusion method at 4°C by mixing 300 nL protein and 200 nL reservoir solution. Obtained crystals were soaked for one day in 0.1 M sodium acetate pH 5.2, 20 mM MgCl2, 10 mM ATP, and 22% (w/v) PEG3350 at 4°C. Before flash-freezing in liquid nitrogen, the crystal was transferred into a cryoprotectant solution consisting of soaking solution supplemented with 33% (v/v) ethylene glycol. Detailed procedures can be found in the *Supporting Information*.

### Hydrogen deuterium exchange mass spectrometry

HDX reactions were performed on the apo enzymes (MBP-*Hs*IP6K1^wt^ or MBP-*Hs*IP6K1^L210V^) or enzymes in the presence of inhibitor (FMP-201300). Prior to initiation of deuterium exchange reactions, enzymes were incubated with inhibitor or DMSO blank for 15 minutes at RT (Final concentration: 3 μM enzyme, 50 μM inhibitor, in a buffer consisting of 20 mM HEPES 7.4, 50 mM NaCl, 1% DMSO). From this mixture, 5 μL was taken per sample and 45 μL of deuterated buffer was added to initiate deuterium exchange (Final concentrations: 300 nM enzyme, 5 μM inhibitor; Final buffer concentration: 20 mM HEPES 7.4, 50 mM NaCl, 85.63% Deuterium, 0.1% DMSO). Reactions were carried out in triplicate at 3 (3s, 30s, 300s) or 4 different time points (0.3s, 3s, 30s, 300s) and were quenched by the addition of 20 μL of ice-cold quench buffer (Final concentration: 0.57 M GuaHCL, 0.86% Formic Acid). Samples were snap-frozen in liquid nitrogen and then stored at -80°C until mass analysis. The peptides that reached significance and met the cutoffs (>6% deuterium incorporation and 0.5 Da with an unpaired student t-test of p<0.05) were plotted on a model composed of the AlphaFold structure of IP6K1 (Q92551), where all affected regions display high (>90%) per-residue confidence scores (pLDDT) and the ATP and InsP6 molecules from the crystal structure of EhIP6KA (PDB: 4O4F).^[53]^ All conclusions are based on the comprehensively described structural similarities between IP6Ks and protein kinases^[25]^, the known crystal structure of the ortholog *Eh*IP6KA^[26b]^, and a homology model for mammalian IP6K2.^[26b]^ Detailed procedures and raw data can be found in the *Supporting Information*.

### Reverse transcription qPCR

The ribosomal DNA transcription assay was performed like described before.^[44]^ Total RNA was extracted using the RNeasy kit from *QIAGEN*. cDNA was generated using *Thermo Fisher Scientific* SuperScript^®^ III Reverse Transcriptase and 2 μg RNA following the vendor’s protocol. Two sets of specific 45S pre-rRNA primers were used as described previously.^[44]^ The qPCR was conducted using SYBR™ Green PCR Master Mix and a StepOnePlus™ RT PCR System. The difference in transcript levels was calculated using the ΔΔct method and technical triplicates. Detailed procedures can be found in the *Supporting Information*.

## Supporting information

HDX Source Data

## Supporting Information

Supporting figures and tables as well as detailed experimental procedures and raw data can be found in the Supporting Information files.

## Acknowledgements

We thank Steven Moss and Kevan Shokat (UCSF) for providing electrophile-sensitive kinase inhibitors. We further thank Stephen Shears and Huanchen Wang (NIEHS) for the *Eh*IP6KA plasmid and suggestions on *Eh*IP6KA expression and crystallization. We thank Janett Tischer from the Protein Production and Characterization Technology Platform at the Max-Delbrück-Center (MDC) for excellent technical assistance and acknowledge the beamline support by the staff of the Helmholtz-Zentrum Berlin für Materialien und Energie at BESSY. We furthermore thank Han Sun and Haoran Liu (FMP) for discussing docking/MD simulation studies with FMP-201300. We are grateful for helpful discussions and comments from all lab members.

## Author contributions

D.F. and T.A. conceived the project. T.A. performed most biochemical experiments, data analysis and wrote the original draft. S.H. optimized the assay for high-throughput screening. C.S performed the high-throughput screens for both wild-type and mutant kinases. M.N. analyzed and curated the raw data from the high-throughput screens. A.S conducted the protein crystallization and structure determination. G.L.D. executed all HDX-MS measurements and performed primary data analysis. D.F., V.H. and J.P.K. supervised the project. All authors reviewed and edited the manuscript.

