## Supporting Information for "An unconventional gatekeeper mutation sensitizes inositol hexakisphosphate kinases to an allosteric inhibitor"

[1]: Leibniz-Forschungsinstitut für Molekulare Pharmakologie, Robert-Rössle-Straße 10, 13125 Berlin, Germany

[2]: Institut für Chemie, Humboldt-Universität zu Berlin, Brook-Taylor-Str. 2, 12489 Berlin, Germany

[3]: Max-Delbrück-Center for Molecular Medicine in the Helmholtz Association (MDC), Technology Platform for Protein Production & Characterization, Robert-Rössle-Straße 10, 13125 Berlin, Germany

* Corresponding author

### Abbreviations

| | |
|---|---|
| 1-B-PP1 | 3-benzyl-1-(tert-butyl)-1*H*-pyrazolo[3,4-*d*]pyrimidin-4-amine |
| 1-NA-PP1 | 1-(tert-butyl)-3-(naphthalen-1-yl)-1*H*-pyrazolo[3,4-*d*]pyrimidin-4-amine |
| 1-NM-PP1 | 1-(tert-butyl)-3-(naphthalen-2-ylmethyl)-1*H*-pyrazolo[3,4-*d*]pyrimidin-4-amine |
| 1PP-Ins$P_5$ | 1-diphosphoinositol 2,3,4,5,6-pentakisphosphate |
| 2,3-dMB-PP1 | 1-(tert-butyl)-3-(2,3-dimethylbenzyl)-1*H*-pyrazolo[3,4-*d*]pyrimidin-4-amine |
| 3-BrB-PP1 | 3-(3-bromobenzyl)-1-(tert-butyl)-1*H*-pyrazolo[3,4-*d*]pyrimidin-4-amine |
| 3-MB-PP1 | 1-(tert-butyl)-3-(3-methylbenzyl)-1*H*-pyrazolo[3,4-*d*]pyrimidin-4-amine |
| 5PP-Ins$P_5$ | 5-diphosphoinositol 1,2,3,4,6-pentakisphosphate |
| 6-MeO-NA-PCA | 5-amino-1-(tert-butyl)-3-(6-methoxynaphthalen-2-yl)-1*H*-pyrazole-4-carboxamide |
| ADP | Adenosine diphosphate |
| AS | Analog-sensitive |
| ATP | Adenosine triphosphate |
| BCA | Bicinchoninic acid |
| BEZ235 | 2-methyl-2-(4-(3-methyl-2-oxo-8-(quinolin-3-yl)-2,3-dihydro-1*H*-imidazo[4,5-*c*]quinolin-1-yl)phenyl)propanenitrile |
| BIRD | Bilinear rotation decoupling |
| BSA | Bovine serum albumin |
| cryo-EM | Cryogenic electron microscopy |
| CV | Column volume |
| DTT | Dithiothreitol |
| *Eh* | *Entamoeba histolytica* |
| gk | gatekeeper |
| HDX-MS | Hydrogen deuterium exchange mass spetrometry |
| HMQC | Heteronuclear single quantum coherence |
| $IC_{50}$ | Half maximal inhibitory concentration |
| IMAC | Immobilized metal ion affinity chromatography |

| | |
|---|---|
| $InsP_6$ | Inositol hexakisphosphate |
| InsPs | Inositol polyphosphates |
| IP6K1 | Inositol hexakisphosphate kinase 1 |
| IP6K2 | Inositol hexakisphosphate kinase 2 |
| IP6K3 | Inositol hexakisphosphate kinase 3 |
| IP6KA | Inositol hexakisphosphate kinase from Entamoeba histolytica |
| IPTG | Isopropyl-β-D-thiogalactopyranosid |
| $k_{cat}$ | Turnover number |
| $K_M$ | Michaelis-Menten constant |
| KO | Knockout |
| LIC | Ligation-independent cloning |
| LOPAC | Library of pharmacologically active compounds |
| MAD | Median absolute deviation |
| MBP | Maltose binding protein |
| NA-PCA | 5-amino-1-(tert-butyl)-3-(naphthalen-2-yl)-1*H*-pyrazole-4-carboxamide |
| NMR | Nuclear magnetic resonance |
| ORF | Open reading frame |
| PAINS | Pan-assay interference |
| PCA | 5-aminopyrazolo-4-carboxamide |
| PCR | Polymerase chain reaction |
| PP1 | 4-amino-1-tert-butyl-3-(4-methylphenyl)-1*H*-pyrazolo[3,4-*d*]pyrimidine |
| PP-InsPs | Inositol pyrophosphates |
| PPIP5K | Inositol hexakisphosphate and diphosphoinositol pentakisphosphate kinase |
| SAR | Structure-activity relationship |
| SD | Standard deviation |
| SDS-PAGE | Sodium dodecyl sulfate polyacrylamide gel electrophoresis |
| TNP | *N*2-(m-trifluorobenzyl)-*N*6-(p-nitrobenzyl)purine |
| WT | Wild type |

### Supporting figures and tables

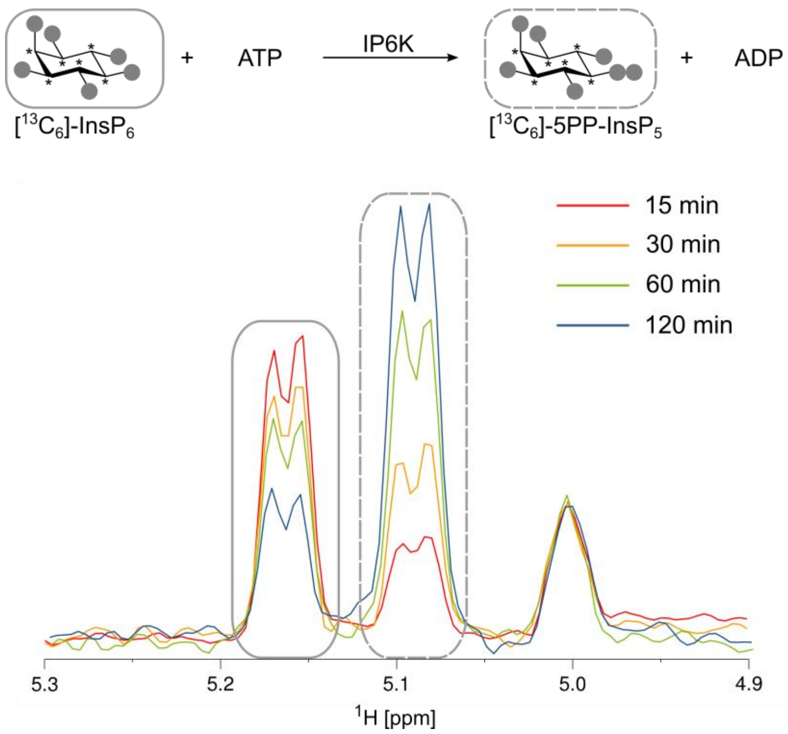

**Figure S1:** Kinase reaction of IP6Ks using fully $^{13}C$-labeled $InsP_6$ as a substrate. The conversion to 5PP-$InsP_5$ was followed by the established spin-echo-difference NMR method.[1] The peaks corresponding to substrate and product are framed with solid or dashed rectangles, respectively.

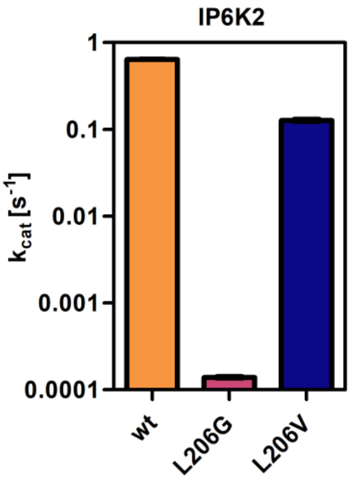

**Figure S2:** Catalytic activities of IP6K2 WT and gatekeeper mutants as indicated by the apparent turnover number. All samples were measured in independent triplicates and error bars represent standard deviation.

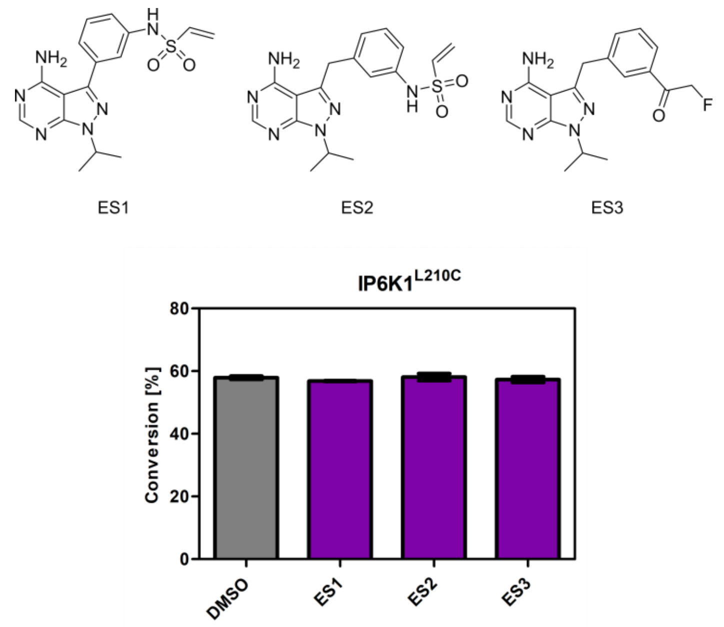

**Figure S3:** Screening of established electrophile-sensitive kinase inhibitors at 10 µM concentration against IP6K1$^{L210C}$ using the NMR assay. All compounds were measured in independent triplicates and error bars represent standard deviation.

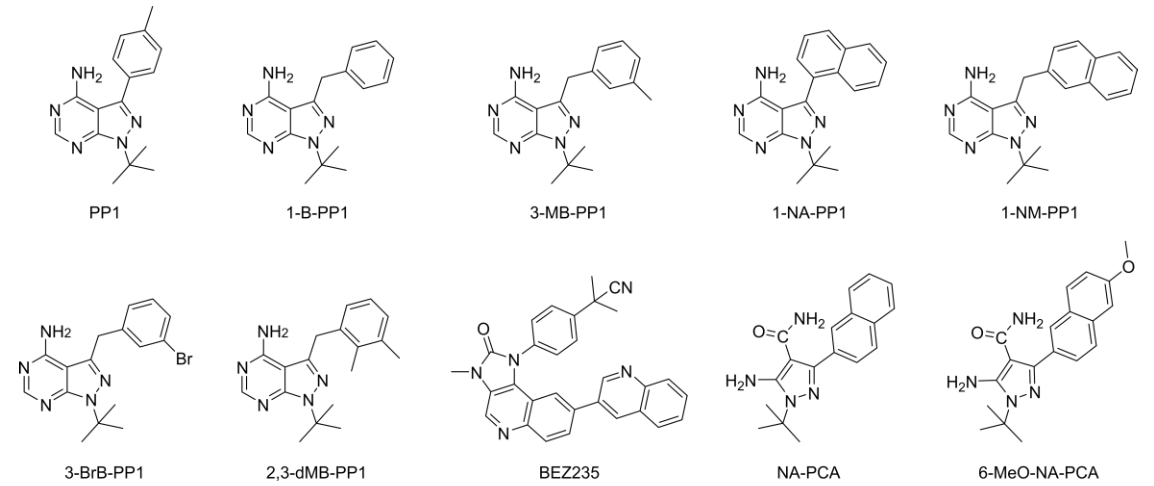

**Figure S4:** Chemical structures of established analog-sensitive kinase inhibitors screened against IP6K1$^{L210V}$.

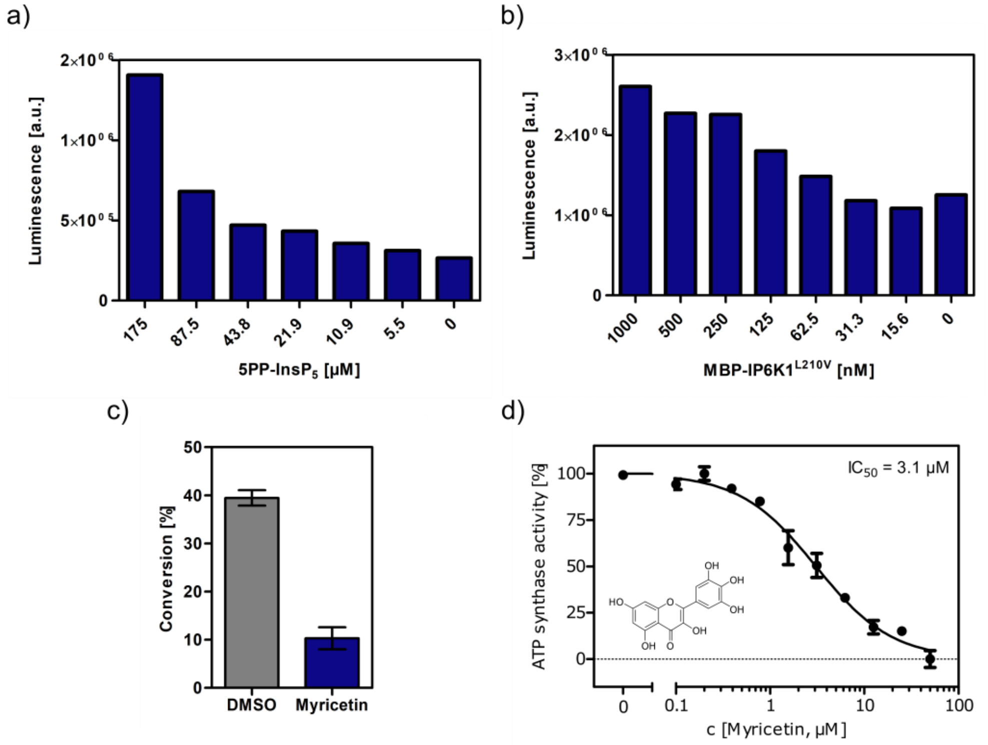

**Figure S5:** Optimization of assay conditions for high-throughput screening. **a) b)** Serial dilution of substrate and protein in single measurements to determine suitable concentrations for the high-throughput screen using the Kinase-Glo® assay. **c)** Substrate conversion at optimized reaction conditions measured by NMR. 10 µM Myricetin was used as positive control. **d)** $IC_{50}$ curve of Myricetin against $IP6K1^{L210V}$ in reverse reaction measured by NMR. All concentrations were measured in independent triplicates and error bars represent standard deviation.

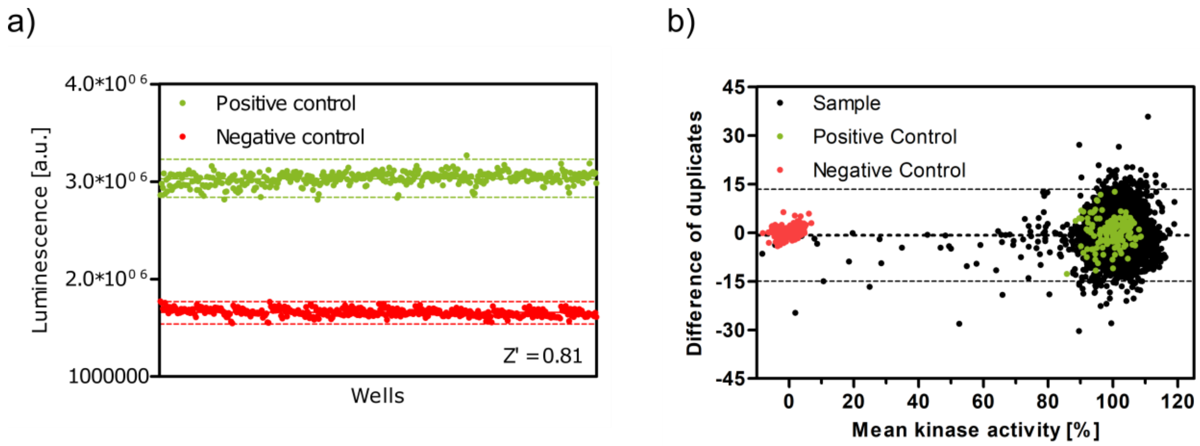

**Figure S6: a)** Z'-plate with positive and negative controls to assess the assay quality and high-throughput screen viability. Dashed lines indicate the ±3*standard deviation values. **b)** Bland-Altman plot of pilot screen. The dotted line indicates the mean of the differences and dashed lines represent the limits of agreement (± 14.2%).

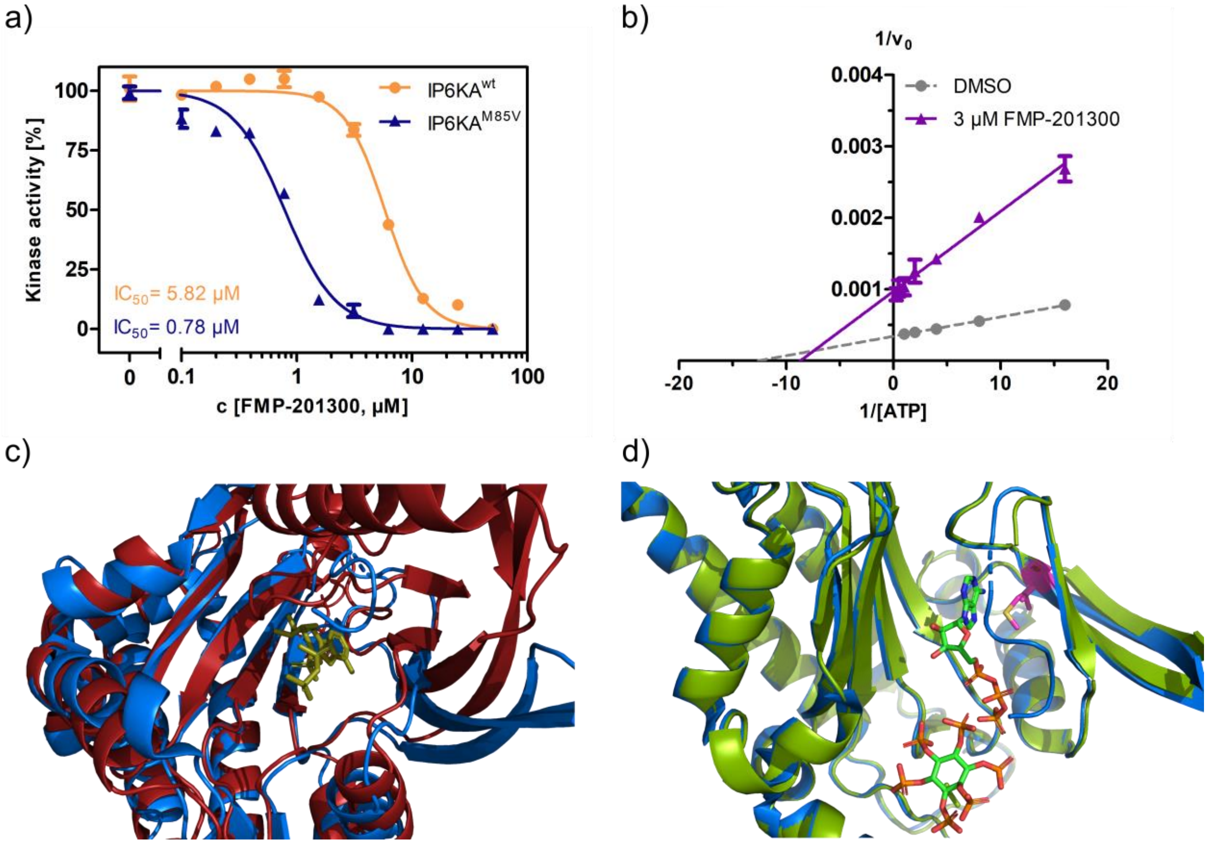

**Figure S7:** Characterization of FMP-201300 binding to *Eh*IP6KA. **a)** $IC_{50}$ curves of FMP-201300 against $IP6KA^{wt}$ and $IP6KA^{M85V}$. **b)** Lineweaver-Burk plot of FMP-201300 against $IP6KA^{wt}$ All points were measured in independent triplicates and error bars represent standard deviation. **c)** Zoom-in on the structural alignment of *Eh*IP6KA (blue) bound to ATP (olive) (PDB: 4O4F) and the AlphaFold structure model of IP6K1 (red) (Q92551).[2] **d)** Zoom-in on the structural alignment of published *Eh*$IP6KA^{wt}$ crystal structure bound to ATP and $InsP_6$ (blue, PDB: 4O4F), and $IP6KA^{M85V}$ crystal structure (green, this study, PDB: 8OMI, bound ATP omitted for clarity). The gatekeeper residue is highlighted in magenta.

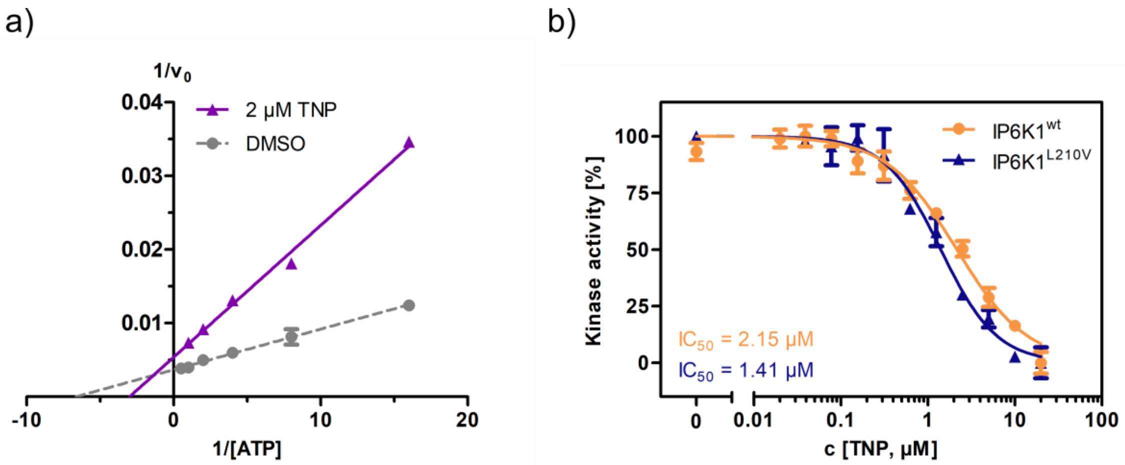

**Figure S8: a)** Lineweaver-Burk plot of TNP against IP6K1$^{wt}$. **b)** $IC_{50}$ curves of TNP against IP6K1$^{wt}$ and IP6K1$^{L210V}$. All points were measured in independent triplicates and error bars represent standard deviation.

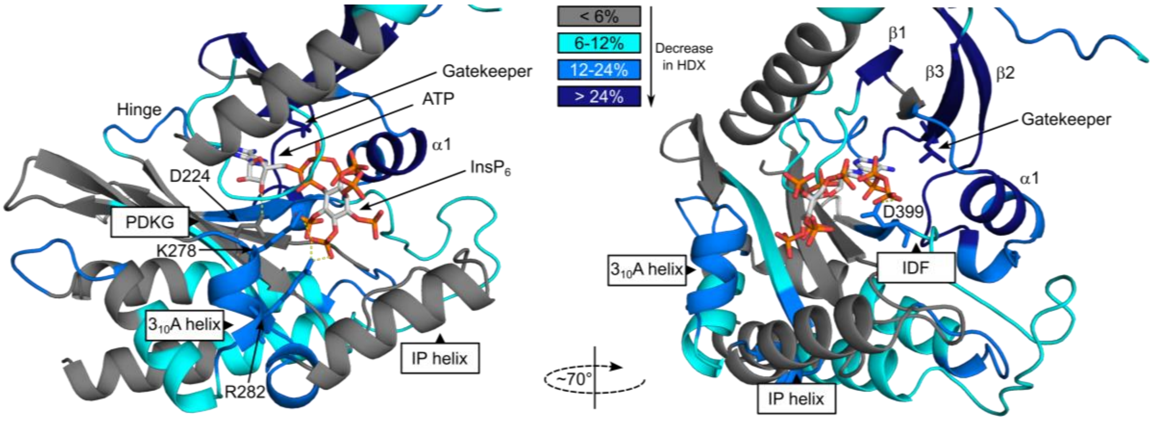

**Figure S9:** HDX-MS shows FMP-201300 binding to IP6K1$^{L210V}$ leads to decreases that correspond to an allosteric mechanism of action. Differences in deuterium exchange rates mapped on a model of IP6K1$^{wt}$ (AlphaFold structure prediction Q92551 with ATP and $InsP_6$ from *Eh*IP6KA docked in the active site (PDB: 4O4F)). Peptides that showed significant differences in HDX and met the cut-offs were included (>6% deuterium incorporation and 0.5 Da with an unpaired student t-test of $p<0.05$). Differences in deuterium incorporation >6%, >12% and >24% compared to the apo enzyme are colored in teal, blue, and navy, respectively. Grey regions had no altered deuterium uptake. Strongly disordered regions and regions of low per-residue confidence scores (pLDDT) were omitted for clarity. Full data can be found in the supporting information.

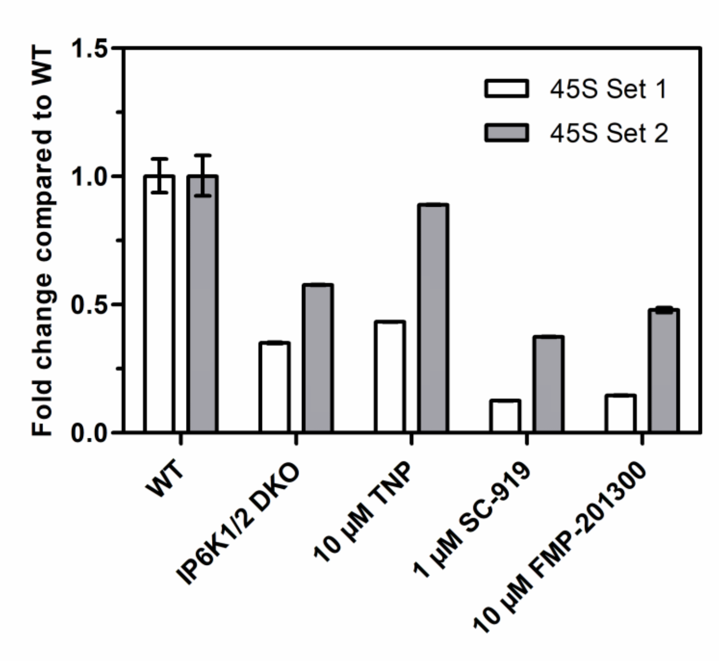

**Figure S10:** RT-qPCR analysis to measure 45S pre-rRNA transcript levels using two different primer sets. Values indicate the fold change in transcript levels in IP6K1/2 double knockout (DKO) cells, or TNP-, SC-919- or FMP-201300-treated HCT116 cells compared to HCT116 WT cells. The fold-change was calculated using the $\Delta\Delta c_t$ method and independent technical triplicates.

**Table S1:** Hit compounds selective for IP6K1$^{L210V}$. FMP-201300, highlighted in green, is the most promising compound as it is not displaying any PAINS motifs, has no known inhibitory activities, and is neither redox-active nor cytotoxic. All values are for IP6K1$^{L210V}$ if not otherwise indicated.

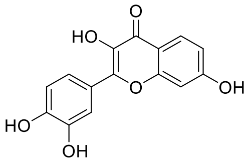

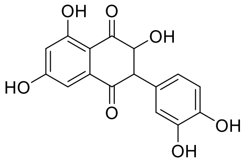

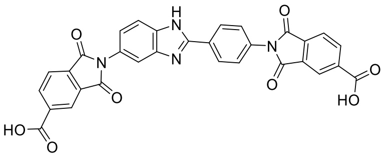

| FMP identifier | 105189 | 200006 | 201300 |
|---|---|---|---|
| Z-score | -18.2 | -18.6 | -24.0 |
| %Activity | 38.3 | 37.4 | 12.5 |
| $IC_{50}$ (IP6K1$^{L210V}$) | 5.13 µM | 4.33 µM | 1.60 µM |
| $IC_{50}$ (IP6K1$^{wt}$) | 85.3 µM | 20.5 µM | 17 µM |

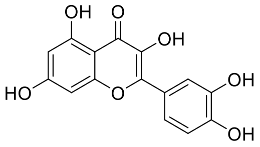

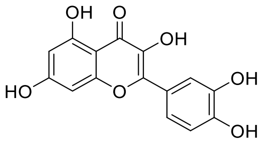

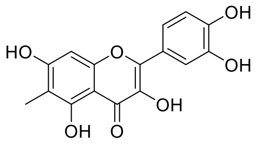

| FMP identifier | 405356 | 710989 | 711656 |
|---|---|---|---|
| Z-score | -14.6 | -17.5 | -24.6 |
| %Activity | 7.8 | 35.6 | 21.3 |
| $IC_{50}$ (IP6K1$^{L210V}$) | 2.86 µM | 3.38 µM | 1.76 µM |
| $IC_{50}$ (IP6K1$^{wt}$) | 12.1 µM | 15.8 µM | 47.5 µM |

**Table S2:** Cytotoxicity of selected hits indicated by cell survival after 72 hours incubation at 10 µM concentration. All compounds were measured in independent triplicates and errors represent standard deviation.

| Compound (10 µM) | Cell line | Cell number (%) |
|---|---|---|
| 105189 | HEK293 | 72.7 ± 9.8 |
| | HepG2 | 67.4 ± 6.4 |
| 200006 | HEK293 | 82.8 ± 9.4 |
| | HepG2 | 104.1 ± 9.9 |
| 201300 | HEK293 | 100.1 ± 4.1 |
| | HepG2 | 99.1 ± 2.8 |
| 405356 | HEK293 | 104.0 ± 1.4 |
| | HepG2 | 65.9 ± 16.9 |

**Table S3:** Redox activity of selected compounds measured *via* a photometric surrogate assay.[3] Reactions contained 50 mM HEPES pH 7.5, 50 mM NaCl, 200 mM DTT, and 20 µM resazurin, and were incubated for one hour at 37°C.

| Compound | Conversion (%) | Z-score |
|---|---|---|
| 105189 | 4.58 | 3.96 |
| 200006 | 2.76 | 1.9 |
| 201300 | -0.0816 | -0.225 |
| 405356 | 0.0343 | 0.674 |

**Table S4:** Data collection and structure refinement statistics.

| | *Entamoeba histolytica* IP6KA$^{M85V}$ variant |
|---|---|
| PDB ID code | 8OMI |
| **Data collection** | |
| Space group | $I4_122$ |
| Cell dimensions | |
| *a*, *b*, *c* (Å) | 102.82, 102.82, 111.61 |
| $\alpha$, $\beta$, $\gamma$ (°) | 90, 90, 90 |
| Resolution (Å) | 37.81-1.77 (1.84-1.77)* |
| $R_{merge}$ (%) | 12.51 (251.1) |
| $< I / \sigma(I) >$ | 8.61 (1.11) |
| Completeness (%) | 99.15 (98.25) |
| Redundancy | 7.5 (7.6) |
| **Refinement** | |
| Resolution (Å) | 1.77 |
| No. unique reflections | 29205 |
| $R_{work}$ / $R_{free}$ (%) | 23.08 / 25.36 |
| No. non-hydrogen atoms | 2233 |
| Protein | 1999 |
| Ligands | 45 |
| Water | 189 |
| Average B-factor (Å$^2$) | |
| Overall | 39.2 |
| Protein | 38.6 |
| Ligands | 49.7 |
| Water | 42.5 |
| R.m.s deviations | |
| Bond lengths (Å) | 0.002 |
| Bond angles (º) | 0.541 |

### Experimental section

#### General information

All reagents were purchased from common suppliers such as Sigma Aldrich, VWR, Roth, TCI, Thermo Scientific or Roche, and used without further purification unless stated otherwise. The entire PCR and LIC equipment (5× Phusion GC buffer, ThermoPol buffer, dNTP's, Phusion DNA polymerase, Taq DNA polymerase, SspI, T4 DNA polymerase) and Dpn1, CutSmart Buffer and SOC medium were purchased from New England Biolabs (NEB). All PP1 inhibitor analogs were acquired from Toronto Research Chemicals and BEZ235 from Cayman Chemical. Deuterated solvents were purchased from Deutero. Luminescence-based detections were performed with Kinase-Glo® assays from Promega.

The pET-His$_6$-MBP-N$_{10}$-TEV LIC cloning vector (2C-T) was a gift from Scott Gradia (Addgene plasmid # 29706 ; http://n2t.net/addgene:29706 ; RRID:Addgene_29706).

PCR was performed on a BIO-RAD C1000 Touch™ Thermal Cycler. Concentrations of DNA and proteins were determined spectroscopically with a Thermo Scientific NANODROP 2000C Spectrophotometer. All recombinant proteins were purified on a BIO-RAD NGC™ chromatography system with a BioFrac Fraction Collector. NMR spectra were recorded on a Bruker SUPERSHIELD™ 600 PLUS. Luminescence assays were read out with a Tecan Infinite M Plex reader. Diffraction data were collected at 100 K at the beamline BL14.1 operated by the Helmholtz-Zentrum Berlin (HZB) in the BESSY II electron storage ring (Berlin-Adlershof, Germany),[4] using a wavelength of 0.9184 Å. Mass spectrometry experiments were performed on an Thermo Scientific Orbitrap Elite.

### Methods

#### Ligation-independent cloning

The LIC cloning vectors were linearized by SspI digest following the manufacturers protocol. After incubation for 30 minutes at 37°C, the linearized plasmids were purified with the QIAquick PCR Purification Kit (50). The IP6K ORFs were amplified by PCR from the pTrc-His constructs using Phusion DNA polymerase and the primers listed in **Table S5**. After initialization at 98°C for one minute, the PCR cycle consisted of denaturation for 10 s at 98°C, annealing for 20 s at 65°C and elongation for 45 s at 72°C, with 30 repetitions. Final elongation was performed for 5 minutes at 72°C. The amplified inserts were purified by PCR purification, DpnI digest and another PCR purification. The complementary overhangs were created by T4 DNA polymerase digest adding dGTP to the linearized plasmid reaction and dCTP to the insert reaction. After incubation for 30 minutes at 22°C, T4 DNA polymerase was heat-inactivated by incubation for 20 minutes at 75°C. Linearized plasmid and IP6K ORF were ligated by combing them in a molar ratio of 1:2 (vector:insert) and incubating them for 30 minutes at room temperature. After addition of 1 µL EDTA (25 mM), 5 µL were used for transformation into *E. coli* TOP10. The successful subcloning was verified by colony PCR from the resulting colonies using Taq DNA polymerase with ThermoPol buffer and the LIC primers listed in **Table S5**. After 8 minutes initialization at 98°C, the PCR cycle consisted of denaturation for 30 s at 95°C, annealing for 30 s at 65°C and elongation for 90 s at 68°C with 30 repetitions. Final elongation was performed for 5 minutes at 68°C.

**Table S5:** Primers used for ligation-independent cloning.

| Primer | Sequence |
|---|---|
| IP6K1 forward | 5’-TACTTCCAATCCAATGCA ATGTGTGTGTGTCAAACCATG-3’ |
| IP6K1 reverse | 5’-TTATCCACTTCCAATGTTATTA TTATTGATTTTCATCGCGCATCTG-3’ |
| IP6K2 forward | 5’-TACTTCCAATCCAATGCA ATGTCTCCGGCGTTTCGTGCTATG-3’ |

| | |
|---|---|
| IP6K2 reverse | 5'-TTATCCACTTCCAATGTTATTA TTATTCGCCGCTTTCTTC-3' |
| IP6K3 forward | 5'-TACTTCCAATCCAATGCA ATGGTCGTTCAAAATTCGGC-3' |
| IP6K3 reverse | 5'-TTATCCACTTCCAATGTTATTA TTCGCCTTCTTGGATATCC-3' |

**PCR mutagenesis**

Plasmid DNA harboring the corresponding IP6K ORF was extracted from an overnight culture of the pET-$His_6$-MBP-$N_{10}$-TEV (*Hs*IP6K) or pET15b-$His_6$ (IP6KA) vector in TB-Amp using a *Qiagen* Miniprep kit. Single point mutations were installed by employing the primers listed in **Table S6**. 50 µL PCR reactions were performed on a *BIO-RAD* C1000 Touch™ Thermal Cycler following the *NEB* Phusion High-Fidelity DNA polymerase protocol using the following temperature program: 5 minutes 98°C → 30 s 98°C → 3 minutes 72°C → Cycle to step 2 30× → 5 minutes 72°C.

**Table S6:** Primers used for PCR mutagenesis to generate gatekeeper mutants of IP6Ks.

| Mutant | | Primer (5'→ 3') |
|---|---|---|
| IP6K1 (L210G) | Forward | CTGCTGGGGGAAAACGTCGTGCATCACTT |
| | Reverse | GTTTTCCCCCAGCAGAAATTTATACAGCT |
| IP6K1 (L210A) | Forward | CTGCTGGCGGAAAACGTCGTGCATCACTT |
| | Reverse | GTTTTCCGCCAGCAGAAATTTATACAGCT |
| IP6K1 (L210V) | Forward | CTGCTGGTGGAAAACGTCGTGCATCACTT |
| | Reverse | GTTTTCCACCAGCAGAAATTTATACAGCT |
| IP6K1 (L210C) | Forward | CTGCTGTGCGAAAACGTCGTGCATCACTT |
| | Reverse | GTTTTCGCACAGCAGAAATTTATACAGCT |
| IP6K2 (L206G) | Forward | ATCCTGGGGGAAAACCTGACCTCCCGCTA |
| | Reverse | GTTTTCCCCCAGGATGAATTTATACTGAT |
| IP6K2 (L206V) | Forward | ATCCTGGTGGAAAACCTGACCTCCCGCTA |
| | Reverse | GTTTTCCACCAGGATGAATTTATACTGAT |
| IP6KA (M85V) | Forward | ATCCGTGTGGAAAACCTGATGTACAAATAC |
| | Reverse | GTTTTCCACACGGATAAATTCCAGTTCAC |

### **Expression and purification of MBP-IP6K constructs**

A 25 mL overnight culture of *E. coli Arctic Express* harboring the IP6K construct on a pET-His$_6$-MBP-N$_{10}$-TEV vector in TB-Amp/Gen was diluted to a final OD$_{600}$ of 0.05 and grown to OD$_{600}$ of 0.7 at 37°C. The temperature was switched to 13°C and expression induced with 0.2 mM IPTG after one hour. After 20h expression, the cells were pelleted by centrifugation (3000 ×g, 10 minutes, 4°C), washed with cold lysis buffer (20 mM Tris-HCl pH 7.4, 150 mM NaCl) and centrifuged again. The pellet was resuspended in 10 mL lysis buffer per gram wet weight and supplemented with lysozyme, DNase I and protease inhibitor. After 30 minutes of incubation on ice, the cells were first homogenized (1× 30s) and then lysed with a microfluidizerTM LM10 at 15.000 psi with three iterations. The cell debris was removed by centrifugation (20.000 ×g, 20 minutes, 4°C) and the supernatant lysate filtered (*VWR* vacuum filter, PES, 0.45 µm). The lysate was loaded onto an equilibrated 5 mL HiTrap IMAC HP column (Ni-NTA, *GE Healthcare*) at a flowrate of 1.5 mL/minute. The column was washed with 10 CV wash buffer (20 mM Tris HCl pH 7.4, 500 mM NaCl, 50 mM imidazole) and the protein eluted with a step gradient of elution buffer (20 mM Tris HCl pH 7.4, 500 mM NaCl, 500 mM imidazole) in wash buffer with 50% B and 100% B over 5 CV, respectively. Protein-containing fractions were pooled and diluted 20-fold with anion exchange start buffer (20 mM Tris HCl pH 8.0). The protein was loaded onto an equilibrated 5 mL HiTrap Q FF (*GE Healthcare*) column at a flowrate of 2 mL/minute, washed with 3 CV of start buffer and eluted with a 0-100% gradient of elution buffer (20 mM Tris-HCl pH 8.0, 1 M NaCl) in start buffer over 10 CV. Protein-containing fractions were concentrated by spin-filtration through 30 kDa cut-off filters to give 1-4 mL and loaded by injection onto a HiLoad 16/60 Superdex 200 pg column equilibrated with 20 mM Tris HCl pH 7.4, 500 mM NaCl, 1 mM DTT. After elution at a flowrate of 0.5 mL/minute, protein-containing fractions were united, concentrated by spin-filtration through 30 kDa cut-off filters, adjusted to 12.5% glycerol, aliquoted and frozen at -80°C. The protein concentration was determined by SDS-PAGE gel electrophoresis with Coomassie staining using a BSA standard.

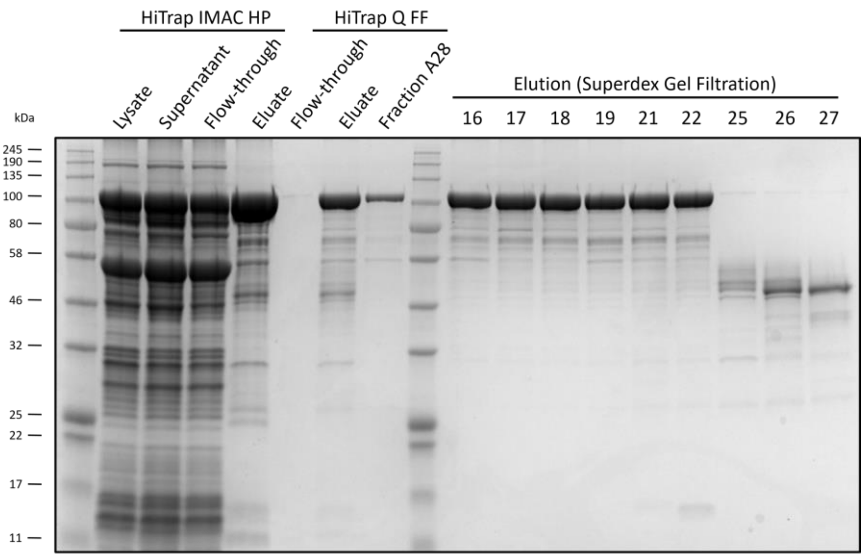

**Figure S11:** Coomassie-stained SDS-PAGE gel from expression of MBP-IP6K1$^{L210V}$.

### Expression and purification of IP6KA constructs

IP6KA was expressed like described before[1] with some slight alterations. An overnight culture of *E. coli Arctic Express* in TB-Amp/Gen harboring the IP6KA construct on a pET15b-His$_6$ vector was diluted 100-fold into 800 mL and grown for six hours at 37°C. The temperature was switched to 13°C and expression induced with 0.1 mM IPTG after one hour. After overnight expression the cells were pelleted by centrifugation (3000 ×g, 10 minutes, 4°C), resuspended in 10 mL lysis buffer (25 mM Tris HCl pH 7.4, 500 mM NaCl, 50 mM imidazole) per gram wet weight and supplemented with lysozyme, DNase I and protease inhibitor. After 30 minutes of incubation on ice the cells were lysed with a microfluidizer™ LM10 at 15.000 psi with five iterations. The cell debris was removed by centrifugation (25000 g, 20 minutes, 4°C) and the supernatant lysate filtered (*VWR* vacuum filter, PES, 0.45 µm). The clarified lysate was loaded onto an equilibrated 5 mL HiTrap IMAC HP (Ni-NTA, *GE Healthcare*) column at a flowrate of 1 mL/minute. The protein was eluted with a gradient of elution buffer (25 mM Tris HCl pH 7.4, 200 mM NaCl, 500 mM imidazole) in lysis buffer from 0-100% over 10 CV. Protein-containing fractions were concentrated by spin filtration through 3.5K cut-off filters and dialyzed overnight against dialysis buffer (20 mM Tris HCl pH 7.4, 200 mM NaCl, 1 mM DTT). The dialyzed protein was adjusted to 25% glycerol, aliquoted and frozen at -80°C. Protein concentration was determined by BCA assay using BSA standards for calibration.

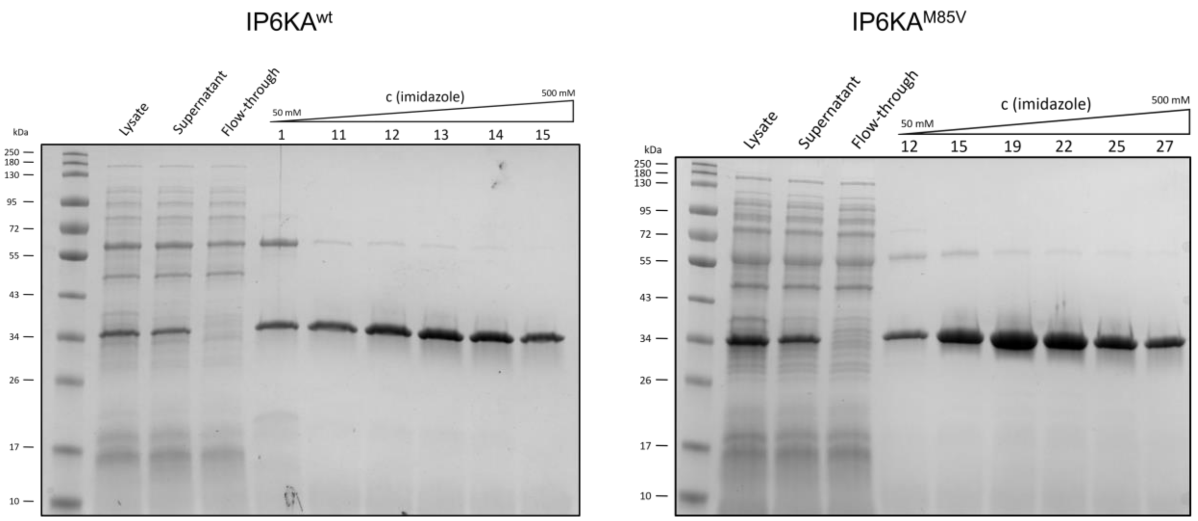

**Figure S12:** Coomassie-stained SDS-PAGE gels from expression of IP6KA$^{wt}$ and IP6KA$^{M85V}$.

### NMR activity assay and $IC_{50}$ measurements for IP6K1 constructs

Enzyme assays were performed in $D_2O$ in a total volume of 150 µL containing 20 mM HEPES pH* 6.8, 50 mM NaCl, 6 mM $MgCl_2$, 1 mM ATP, 0.2 mg/mL BSA, 1 mM DTT, 5 mM creatine phosphate and 1 U/mL creatine kinase (ATP-regenerating system) and 50 nM kinase if not otherwise stated. HEPES, NaCl and BSA were prepared as buffer A and DTT and creatine phosphate as buffer B in $D_2O$ and were adjusted to pH* 6.8, respectively. If applicable, inhibitor was added as a DMSO-$d_6$ stock in a two-fold dilution series at a final DMSO concentration of 1% if not otherwise mentioned. Otherwise, DMSO-$d_6$ was added to the same percentage. The samples were equilibrated at 37°C for 3 minutes and the reaction initiated by adding 100 µM [$^{13}C_6$]-$InsP_6$. The reaction was quenched with 400 µL quenching solution (20 mM EDTA pD 6.0, 68.75 mM NaCl) and transferred into an NMR tube before being measured on a *Bruker* SUPERSHIELD™ 600 PLUS with an NMR pulse program developed in our group.[1] $IC_{50}$ curves and values were calculated in *GraphPad* Prism (Version 5.04.) using nonlinear regression, where the DMSO control defines 0% inhibition and a no kinase control defines 100% inhibition.

### NMR activity assays and $IC_{50}$ measurements for IP6KA constructs

Enzyme assays were performed in $D_2O$ in a total volume of 150 µL containing 20 mM MES pH* 6.4, 50 mM NaCl, 6 mM $MgCl_2$, 1 mM ATP, 0.2 mg/mL BSA, 1 mM DTT,

5 mM creatine phosphate and 1 U/mL creatine kinase (ATP-regenerating system) and 50 nM kinase if not otherwise stated. MES, NaCl and BSA were prepared as buffer A and DTT and creatine phosphate as buffer B in $D_2O$ and were adjusted to pH* 6.4, respectively. If applicable, inhibitor was added as a DMSO-$d_6$ stock in a two-fold dilution series at a final DMSO concentration of 1% if not otherwise mentioned. Otherwise, DMSO-$d_6$ was added to the same percentage. The samples were equilibrated at 37°C for 3 minutes and the reaction initiated by adding 100 µM [$^{13}C_6$]-$InsP_6$. The reaction was quenched with 400 µL quenching solution (20 mM EDTA pD 6.0, 68.75 mM NaCl) and transferred into an NMR tube before being measured on a *Bruker* SUPERSHIELD™ 600 PLUS with an NMR pulse program developed in our group.[1] $IC_{50}$ curves and values were calculated in *GraphPad* Prism (Version 5.04.) using nonlinear regression, where the DMSO control defines 0% inhibition and a no kinase control defines 100% inhibition.

**Kinase-Glo® assay optimization**

The following solutions were prepared as 4× stock solutions: kinase buffer (HEPES pH 7.4, NaCl, BSA, $MgCl_2$, Tween-20, DTT and IP6K1$^{L210V}$), DMSO/inhibitor, ADP and 5PP-$InsP_5$. Final concentrations after optimization were 1 mM ADP, 20 mM HEPES, 50 mM NaCl, 0.2 mg/mL BSA, 2 mM $MgCl_2$, 0.05% Tween-20, 1 mM DTT, 250 nM IP6K1$^{L210V}$ and 50 µM 5PP-$InsP_5$. The reactions were carried out in 384-well plates following this procedure. 5 µL 4× kinase buffer (buffer only for no kinase controls), 5 µL 4× DMSO or inhibitor and 5 µL 4× ADP solution were subsequently added to all wells. After 10 minutes equilibration, 5 µL 4× 5PP-$InsP_5$ were added to start the reaction (milli-Q water for negative controls). After three hours at room temperature, 20 µL of *Promega* Kinase-Glo Plus® reagent were added and the luminescence read out with a *Tecan* Infinite M Plex reader using 100 ms exposure time after 10 minutes of equilibration.

**Z'-plate measurement and high-throughput screen**

ADP was prepared in a 2× stock solution in milli-Q water and the pH adjusted to 7.0. The following solutions were prepared as 4× stock solutions: kinase buffer (HEPES pH 7.4, NaCl, BSA, $MgCl_2$, Tween-20, DTT and IP6K1$^{L210V}$) and 5PP-$InsP_5$. Final concentrations were 1 mM ADP, 20 mM HEPES, 50 mM NaCl, 0.2 mg/mL BSA, 2 mM

$MgCl_2$, 0.05% Tween-20, 1 mM DTT, 250 nM IP6K1$^{L210V}$ and 50 µM 5PP-$InsP_5$. The reactions were carried out in 384-well plates following this procedure. 10 µL 2× ADP solution were added to all wells, then 0.2 µL DMSO or compound (1 mM in DMSO) were added followed by the addition of 5 µL 4× kinase buffer. For dose-response curves, compounds were added in serial dilution (2-fold serial dilutions in DMSO across multiple plates prior to compound transfer to assay plates, 9 concentrations in total). After 10 minutes equilibration, 5 µL 4× 5PP-$InsP_5$ were added to start the reaction (milli-Q water for negative controls). After three hours at room temperature, 20 µL of *Promega* Kinase-Glo Plus® reagent were added and the luminescence read out with a *Tecan* Infinite M Plex or *Perkin Elmer* EnVision reader using 100 ms exposure time after 10 minutes of equilibration. In the counter screen, dose-response curves were measured in the absence of kinase and presence of 50 µM ATP.

**Michaelis-Menten and Lineweaver-Burk kinetics**

Kinetic assays were performed in $D_2O$ in a total volume of 500 µL containing the same buffer components for IP6K and IP6KA constructs like stated for the NMR activity assays above. However, only 5 mM $MgCl_2$ were added as a constant amount while ATP was added as ATP*Mg solution prepared in two-fold dilution starting from 50 mM. ATP*Mg was added to final ATP concentrations ranging from 4 mM to 62.6 µM. If applicable, inhibitor was added as a DMSO-$d_6$ stock at a final DMSO concentration of 1%. Otherwise, DMSO-$d_6$ was added to the same percentage. The samples were equilibrated at 37°C for 5 minutes and the reaction initiated by adding 100 µM [$^{13}C_6$]-$InsP_6$. The reaction time was adjusted so that conversion would not exceed 20% to remain in the linear range for initial velocity measurements. The reaction was quenched with 38 µL 700 mM EDTA pH* 8.0 and transferred into an NMR tube before being measured on a *Bruker* SUPERSHIELD™ 600 PLUS with an NMR pulse program developed in our group.[1] Initial velocities were calculated by dividing the amount of generated product by the product of reaction time in seconds and mass of the kinase in mg. $K_M$ and Lineweaver-Burk plots were generated in GraphPad Prism (Version 5.04.) using non-linear and linear regression, respectively.

**Protein crystallization and structure determination**

The protein construct for crystallization was expressed as follows. An overnight culture of *E. coli Arctic Express* (DE3) harboring pDest-566-MBP-IP6KA$^{M85V}$ (27-270) was diluted to a final OD600 of 0.01 in 800 mL TB and grown for 5.5 hours at 37°C. The temperature was switched to 13°C and expression induced with 0.1 mM IPTG after one hour. After 20h expression, the cells were pelleted by centrifugation (3000 ×g, 10 minutes, 4°C) and frozen at -80°C. A week later, the pellet (10 g) was resuspended in 100 mL lysis buffer (50 mM Tris HCl pH 7.4, 500 mM NaCl, 1 mM $MgCl_2$) supplemented with lysozyme, DNase I and protease inhibitor. After 30 minutes of incubation on ice, the cells were homogenized for 30 s and lysed with a microfluidizerTM LM10 at 15.000 psi with three iterations. The cell debris was removed by centrifugation (20.000 ×g, 20 minutes, 4°C) and the supernatant lysate filtered (VWR vacuum filter, PES, 0.45 µm). The lysate was adjusted to 50 mM imidazole and loaded onto an equilibrated 5 mL HiTrap IMAC HP column (*GE Healthcare*) at a flowrate of 1.5 mL/minute. The column was washed thoroughly with wash buffer (25 mM Tris HCl pH 7.4, 500 mM NaCl, 50 mM imidazole) and the protein eluted with a 0-100% gradient of elution buffer (25 mM Tris HCl pH 7.4, 500 mM NaCl, 500 mM imidazole) in wash buffer over 10 CV. Protein-containing fractions were pooled, adjusted to 1 mM β-mercaptoethanol and 0.5 mg TEV protease from (a gift from Martina Leidert) and dialyzed in a 8 kDa cut-off tube overnight against dialysis buffer (50 mM Tris HCl pH 7.4, 200 mM NaCl, 1 mM DTT). Then, the volume was reduced to 2 mL by spin-filtration through 10 kDa cut-off filters and the TEV-mixture was loaded by injection onto two connected 5 mL MBPTrap HP (GE Healthcare) columns at a flow rate of 2 mL/minute. The columns were washed with 3 CV loading buffer (25 mM Tris-HCl pH 7.4, 150 mM NaCl, 1 mM DTT) at a flowrate of 2 mL/minute and MBP-proteins eluted with 3 CV of elution buffer (25 mM Tris-HCl pH 7.4, 150 mM NaCl, 1 mM DTT, 10 mM D-maltose) at a flowrate of 3 mL/minute. Protein-containing fractions from the flow-through were concentrated by spin-filtration through 3 kDa cut-off filters to give 3 mL and again loaded onto two connected 5 mL MBPTrap HP columns at a flow rate of 2 mL/minute. The columns were washed with 3 CV loading buffer at a flowrate of 2 mL/minute and MBP eluted with 3 CV of elution buffer at a flowrate of 3 mL/minute. Protein-containing fractions from the flow-through were concentrated by spin-filtration through 3 kDa cut-off filters to give 2.5 mL and loaded by injection onto a HiLoad 16/60 Superdex 75 pg column equilibrated with 20 mM Tris-HCl pH 7.4, 150 mM NaCl and 1 mM DTT. After elution at a flowrate of 0.2 mL/minute, protein-containing fractions were concentrated by spin-filtration through 10 kDa cut-off filters to give 300 µL. The concentration of the protein was determined by absorbance spectroscopy at 280 nm using a calculated extinction coefficient of 35870 M-1 cm-1. 800 mL culture yielded 11 mg pure protein (17 mg/mL) that was aliquoted to 50 µL aliquots and frozen at -80°C.

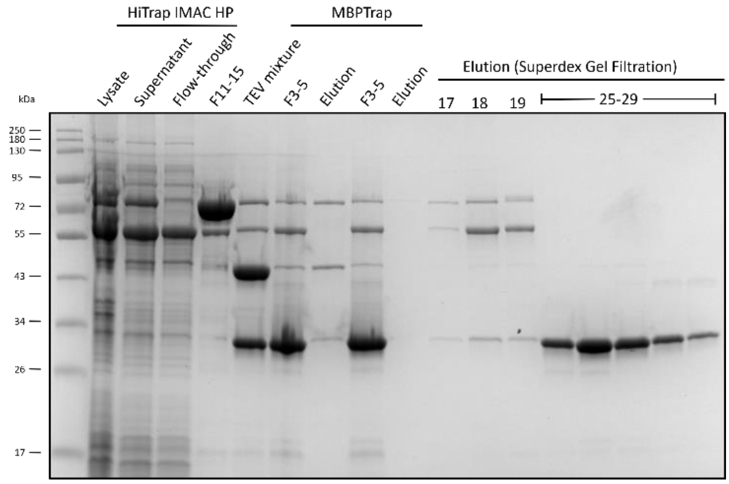

**Figure S13:** Coomassie-stained SDS-PAGE gel from expression of IP6KA$^{M85V}$ (27-270).

The purified *Entamoeba histolytica* IP6KA$^{M85V}$ (27-270) variant protein (17 mg/mL in 20 mM Tris-HCl pH 7.4, 150 mM NaCl, 1 mM DTT) was complexed with 10 mM $MgCl_2$ and 10 mM ATP, and crystallized using the sitting-drop vapor-diffusion method at 4°C by mixing 300 nL protein and 200 nL reservoir solution (1 mM $MgCl_2$, 460 mM $NaH_2PO_4$). Obtained crystals were soaked for one day in 0.1 M sodium acetate pH 5.2, 20 mM $MgCl_2$, 10 mM ATP, and 22% (w/v) PEG3350 at 4°C. Before flash-freezing in liquid nitrogen, the crystal was transferred into a cryoprotectant solution consisting of soaking solution supplemented with 33% (v/v) ethylene glycol. Diffraction data were collected at 100 K at the beamline BL14.1 operated by the Helmholtz-Zentrum Berlin (HZB) in the BESSY II electron storage ring (Berlin-Adlershof, Germany),[4] using a wavelength of 0.9184 Å. Data were processed with the program XDSAPP.[5] The structure was solved by molecular replacement using the program PHASER[6] and the known wild-type *Entamoeba histolytica* IP6KA crystal structure with PDB ID code 4o4f as search model.[7] The structure was refined using PHENIX[8] and the graphics program COOT was used for model building and visualization.[9] Figures were created with PYMOL.[10]

**Hydrogen deuterium exchange mass spectrometry**

Deuterium Exchange Reactions

HDX reactions were performed on the apo enzymes (MBP-*Hs*IP6K1$^{wt}$ or MBP-*Hs*IP6K1$^{L210V}$) or enzymes in the presence of inhibitor (FMP-201300). Prior to initiation of deuterium exchange reactions, enzymes were incubated with inhibitor or DMSO blank for 15 minutes at RT (Final concentration: 3 µM enzyme, 50 µM inhibitor, in a buffer consisting of 20 mM HEPES 7.4, 50 mM NaCl, 1% DMSO). From this mixture, 5 µL was taken per sample and 45 µL of deuterated buffer was added to initiate deuterium exchange (Final concentrations: 300 nM enzyme, 5 uM inhibitor; Final buffer concentration: 20 mM HEPES 7.4, 50 mM NaCl, 85.63% Deuterium, 0.1% DMSO). Reactions were carried out in triplicate at 3 (3s, 30s, 300s) or 4 different time points (0.3s, 3s, 30s, 300s) and were quenched by the addition of 20 µL of ice-cold quench buffer (Final concentration: 0.57 M GuaHCL, 0.86% Formic Acid). Samples were snap-frozen in liquid nitrogen and then stored at -80°C until mass analysis.

Peptide Digestion, Identification, and Measurement of Deuterium Incorporation

Samples were rapidly thawed and injected onto an ultra-performance liquid chromatography (UPLC) at 2°C. The protein was run over an immobilized pepsin column (Affipro, product number: AP-PC-001), and the peptides were collected onto a pre-column trap (Hypersil GOLD, 2.1 mm x 10 mm, *Thermo Fischer*, product number: 25005-012101). The trap was eluted in line with a an Acquity UPLC BEH C18 column (*Waters*, 130Å, 1.7 um, 2.1 mm x 100 mm, product number:186002352) with a gradient of 10-43% buffer B over 18 minutes (buffer A 0.1% formic acid, buffer B 99.9% acetonitrile, 0.1% formic acid). Mass spectrometry experiments were performed on an Orbitrap Elite (*Thermo Scientific*) acquiring over a mass range from 308-2000 m/z using an electrospray ionization source operated at a temperature of 22°C and a spray voltage of 3.8 kV. Peptides were identified using data-dependent acquisition methods following tandem MS/MS experiments. MS/MS datasets were analyzed using MaxQuant against a database of purified proteins and known contaminants with an FDR cutoff of 1%.

Mass Analysis of Peptide Centroids

Deuterium incorporation of peptides was automatically calculated using HDExaminer Software (Sierra Analytics). Peptides were manually inspected for correct parameters (charge state, retention time, isotopic distribution etc). Results are presented as relative levels of deuterium incorporation with the only control for back exchange as the deuterium levels in the buffer. Changes in any peptide at any time point greater than specified cut-offs (>6% deuterium incorporation and 0.5 Da between apo and different conditions with an unpaired student t-test of $p<0.05$) were considered significant.

Interpretation of results

The results were plotted on a model composed of the AlphaFold structure model of IP6K1 (Q92551), where all affected regions display high (>90%) per-residue confidence scores (pLDDT).[2] For orientation, ATP and $InsP_6$ molecules from the crystal structure of *Eh*IP6KA (PDB: 4O4F) were docked in the active site. All conclusions are based on the comprehensively described structural similarities between IP6Ks and protein kinases[11], the known crystal structure of the ortholog *Eh*IP6KA[7], and a homology model for mammalian IP6K2.[7]

**Reverse transcription qPCR**

The ribosomal DNA transcription assay was performed like described before.[12] HCT116 cells were grown to 90-95% confluency before being treated with inhibitor or DMSO for 5 hours (0.1% final DMSO concentration). Total RNA was extracted using the RNeasy kit from *QIAGEN*. cDNA was generated using *Thermo Fisher Scientific* SuperScript® III Reverse Transcriptase and 2 µg RNA following the vendor's protocol. Two sets of specific 45S pre-rRNA primers were used as described previously.[12] The cDNA was diluted 10-fold before conducting qPCR using SYBR™ Green PCR Master Mix and a StepOnePlus™ RT PCR System. The difference in transcript levels was calculated using the $\Delta\Delta c_t$ method and technical triplicates.
